# XPC deficiency increases risk of hematologic malignancies through mutator phenotype and characteristic mutational signature

**DOI:** 10.1101/2020.07.13.200667

**Authors:** Andrey A. Yurchenko, Ismael Padioleau, Bakhyt T. Matkarimov, Jean Soulier, Alain Sarasin, Sergey Nikolaev

## Abstract

Recent studies demonstrated a dramatically increased risk of leukemia in patients with a rare genetic disorder, Xeroderma Pigmentosum group C (XP-C), characterized by constitutive deficiency of global genome nucleotide excision repair (GG-NER). However, the genetic mechanisms of non-skin cancers in XP-C patients remain unexplored. In this study, we analyzed a unique collection of internal XP-C tumor genomes including 6 leukemias and 2 sarcomas. We observed an average of 25-fold increase of mutation rates in XP-C vs. sporadic leukemia which we presume leads to its elevated incidence and early appearance. In all XP-C tumors predominant mutational process is characterized by a distinct mutational signature, highlighting a specific mutational pattern in the context of GG-NER deficiency. We observed a strong mutational asymmetry with respect to transcription and the direction of replication in XP-C tumors suggesting association of mutagenesis with bulky purine DNA lesions of probably endogenous origin. These findings suggest existence of a balance between formation and repair of bulky DNA lesions by GG-NER in human body cells which is disrupted in XP-C patients leading to internal cancers.

## INTRODUCTION

Xeroderma Pigmentosum is a group of rare recessive genetic disorders which includes seven complementation groups (A-G) characterized by constitutive biallelic deficiency of Nucleotide Excision Repair (NER) pathway, and XP variant (loss of polymerase η; Lehmann et al., 2011). NER serves as a primary pathway for repairing various helix-distorting DNA adducts. The NER is subdivided into global genome (GG-NER) and transcription-coupled (TC-NER) sub-pathways which preferentially operate genome-wide and on the transcribed DNA strand of genes respectively. XP patients demonstrate striking tumor-prone phenotype with near 10000-times increased risk of non-melanoma skin cancer and 2000-times risk of melanoma due to the inability of cells to efficiently repair the major UV photoproducts (Bradford et al., 2011; Kraemer et al., 1994). XP complementation group C (XP-C) characterized by GG-NER deficiency (but with an unaffected TC-NER) is one of the most tumor susceptible subtypes of XP (Sethi et al., 2013). Moreover, it was hypothesized that XP patients may harbor 10-20 times increased risk to some types of internal tumors including leukemia, sarcomas (Kraemer, 1987) and thyroid nodules (Hadj-Rabia et al., 2013; Jerbi et al., 2016).

Two recent studies reported a more than a thousand-fold increased risk of hematological malignancies in independent cohorts of XP-C patients (Oetjen et al., 2019; Sarasin et al., 2019) which demonstrated mainly myelodysplastic syndrome with secondary acute myeloid leukemia manifestation. The genetic mechanism of increased risk of internal tumors in XP patients is not well understood.

Experiments with animal XP-C models demonstrated high incidence of liver and lung cancer (Melis et al., 2008) as well as 30-fold increase of spontaneous mutation rate in *Hprt* gene in T-lymphocytes of 1 year old mice (Wijnhoven et al., 2000). Induction of oxidative stress has been shown to further increase the somatic mutagenesis in *Xpc^−/−^*-deficient mice with steady accumulation with age (Melis et al., 2013). A similar tumor-prone phenotype was observed in *Ddb2*/*Xpe* deficient mice with impaired GG-NER pathway: these animals developed broad spectrum of tumors with particularly high incidence of hematopoietic neoplasms (Yoon et al., 2005).

In this work we performed whole genome sequencing (WGS) of a unique collection of internal tumors from XP-C patients to demonstrate that the constitutive GG-NER deficiency causes mutator phenotype rendering susceptibility to hematological malignancies. A particular genomic mutational signature explains the majority of mutations in the studied XP-C leukemia and sarcomas. Observed mutational profiles indicate that mutational process is associated with lesions formed from purine bases. This is the first work which explores mutational patterns in XP-C patients beyond cutaneous malignancies genome-wide.

## RESULTS

We sequenced whole genomes of 6 myeloid leukemia, 1 uterine rhabdomyosarcoma and 1 breast sarcoma along with paired normal tissues from unrelated patients, representing XP-C, the most frequent group of XP in the Northern Africa and Europe (Soufir et al., 2010) and created a catalogue of 202467 somatic mutations (Table 1). Seven out of eight samples harbored a founder c.1643_1644 delTG mutation characteristic of given XP-C population (Soufir et al., 2010) (Table 1). The patients developed internal tumors early in life, between 12 and 30 years of age (median age of tumor diagnosis - 24 years, Table 1). XP-C cancers contained somatic copy number aberrations (SCNAs) and mutations which are characteristic for corresponding types of sporadic malignancies: mutations in *TP53* and deletions of chromosomes 5 and 7 in leukemia, biallelic loss of *CDKN2A* in breast cancer and highly unstable genome of rhabdomyosarcoma (Supplementary Table 1).

**Table 1.** Description of the studied XP-C tumors. a: delTG refers to c.1643_1644 delTG; p.Val548AlafsX572 (Soufir et al., 2010), IVS12 refers to the splice site mutation NM_004628:exon13:c.2251-1G>C (Cartault et al., 2011). b: RAEB - Refractory Anemia with Excess Blasts, AML - Acute Myeloid Leukemia, MDS - Myelodysplastic Syndrome, T-ALL - T-cell Acute Lymphoblastic Leukemia. *Tumor samples used for genomic sequencing.

| Sample | Geographic<br>familial<br>origin | Homozygous<br>*XPC* gene<br>mutation<sup>a</sup> | Sex | Age at<br>diagnosis | Diagnosis<sup>b</sup> | SBS | ID | DBS |
| --- | --- | --- | --- | --- | --- | --- | --- | --- |
| SA009T1 | Spain | delTG | F | 24 | RAEB-2, AML*<br>with MDS-related<br>changes | 36745 | 3147 | 235 |
| SA002T2 | Algeria | delTG | F | 14 | Uterine<br>Rhabdomyosarcoma* | 5692 | 280 | 43 |
| SA006T2 | Algeria | delTG | M | 29 | AML | 33230 | 2322 | 198 |
| SA008T6 | Tunisia | delTG | F | 24 | RAEB-1, RAEB-2,<br>AML-6* | 37783 | 2377 | 245 |
| SA012T2 | Morocco | delTG | F | 29 | AML-6 | 33821 | 2575 | 160 |
| SA007T3 | Comorian<br>Archipelago | IVS12 | F | 30 | Breast sarcoma* | 4787 | 451 | 34 |
| SA010T2 | Tunisia | delTG | M | 16 | AML-6 | 17685 | 1722 | 99 |
| SA011T2 | Morocco | delTG | M | 12 | T-ALL, RAEB-1,<br>AML* | 17274 | 1464 | 98 |

We identified 14.5-31.2 (mean 24.6) fold increase in the number of somatic mutations in XP-C leukemia samples relative to the sporadic myeloid neoplasms (Mann–Whitney U test, two-sided, *P* = 5.8e-05) and the absence of such an effect for XP-C sarcomas (Fig. 1a). This effect was consistent for single base substitutions (SBS), small indels (ID) and double base substitutions (DBS, Fig 1a).

**Fig. 1.**
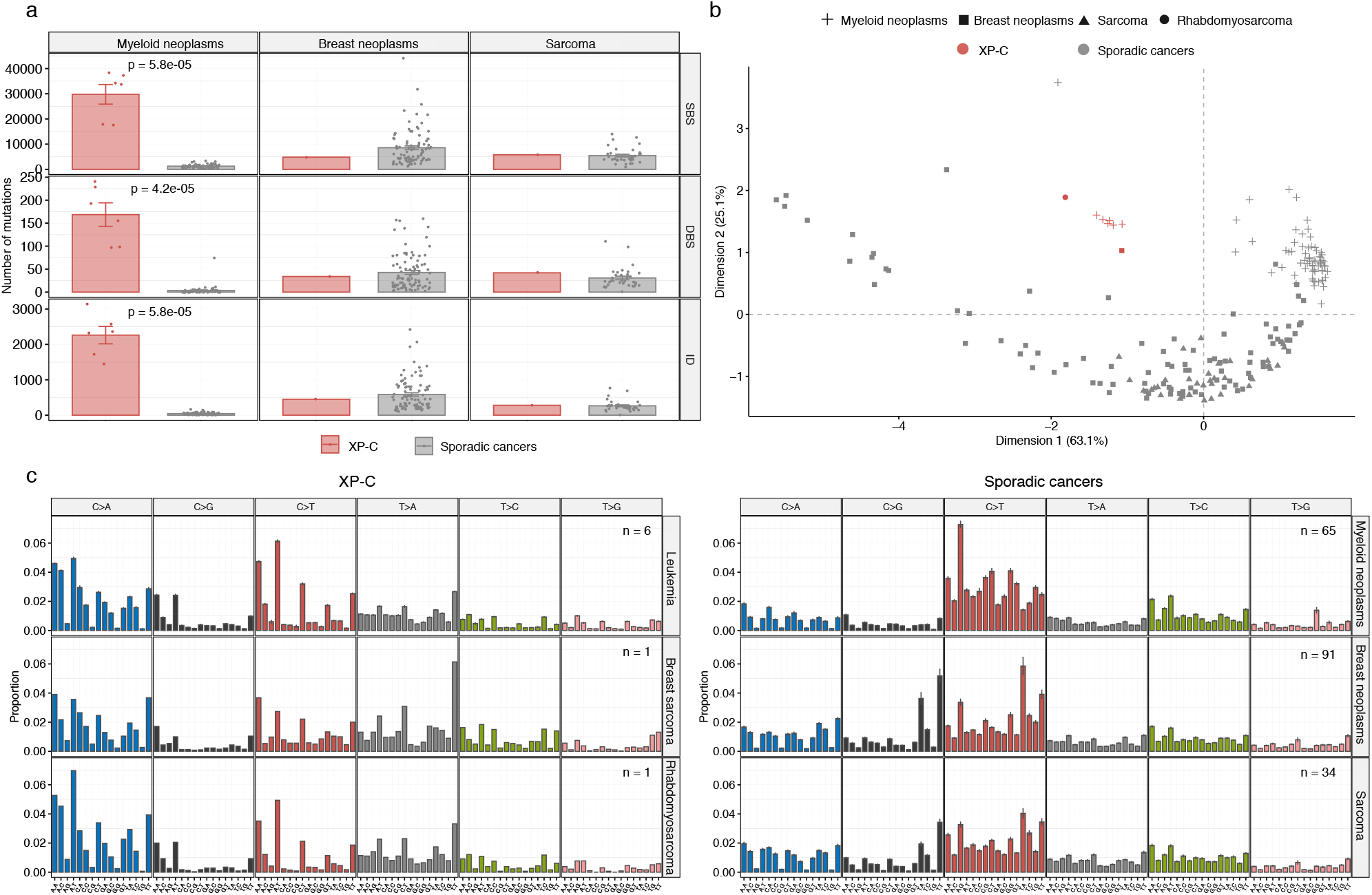
Mutational load and profiles of XP-C and 190 tissue-matched sporadic cancers. **a**, Number of SBS (single base substitutions), DBS (double base substitutions) and ID (indels) in XP-C and sporadic cancers with indicated SEM intervals. The difference is highly significant for myeloid neoplasms (Mann–Whitney U test, two-sided), but number of mutations in single samples of breast sarcoma and rhabdomyosarcoma are in the range of sporadic tumors. **b**, Multidimension scaling plot based on the Cosine similarity distance between the mutational profiles of the samples. XP-C tumors clearly groups together and are distant from tissue-matched sporadic cancers. **c**, Trinucleotide-context mutational profiles (SEM intervals are shown in case of multiple samples). X-axis represents the nucleotides upstream and downstream of mutation. XP-C tumors demonstrate high similarity with each other (left panel), but profiles of sporadic cancers (right panel) are different from them.

The genomic mutational profiles in XP-C tumors were similar between each other irrespectively of the tumor type (average pairwise Cosine similarity of 0.964 (from 0.886 to 0.998)) (Figs. 1b, c;Supplementary Fig. 1, Supplementary Tables 3,4,5) but were different from tissue-matched sporadic tumors (Fig 1b,c). The distinct grouping of XP-C tumors based on mutational profiles was further confirmed in the context of 190 sporadic tissue-matched cancers by Multidimension Scaling analysis (Fig 1b). The mutational patterns of indels was dominated by single nucleotide deletions of C:G and T:A bases in homopolymer stretches and dinucleotide deletions in repeats (Supplementary Fig. 1b). The dinucleotide substitutions were not overrepresented by specific classes and demonstrated a broad range of contexts (Supplementary Fig. 1c).

To better understand mutational processes operating in XP-C cancers we extracted mutational signatures from XP-C and sporadic tissue-matched tumors with the non-negative matrix factorization approach (Alexandrov et al., 2013a) (NMF). Seven signatures were extracted from this dataset (Supplementary Figs. 2a,b) and one of them, Signature “C” explained on average 83.1% of mutations in the XP-C samples (57% in breast sarcoma, 88.9% in rhabdomyosarcoma and 84.1-88.7% in leukemia) while in sporadic tumors only small contribution (average 9.7%, range 0-34.3%) of signature “C” was observed (Figs. 2a,b., Supplementary Figs. 2c,d).

**Fig. 2.**
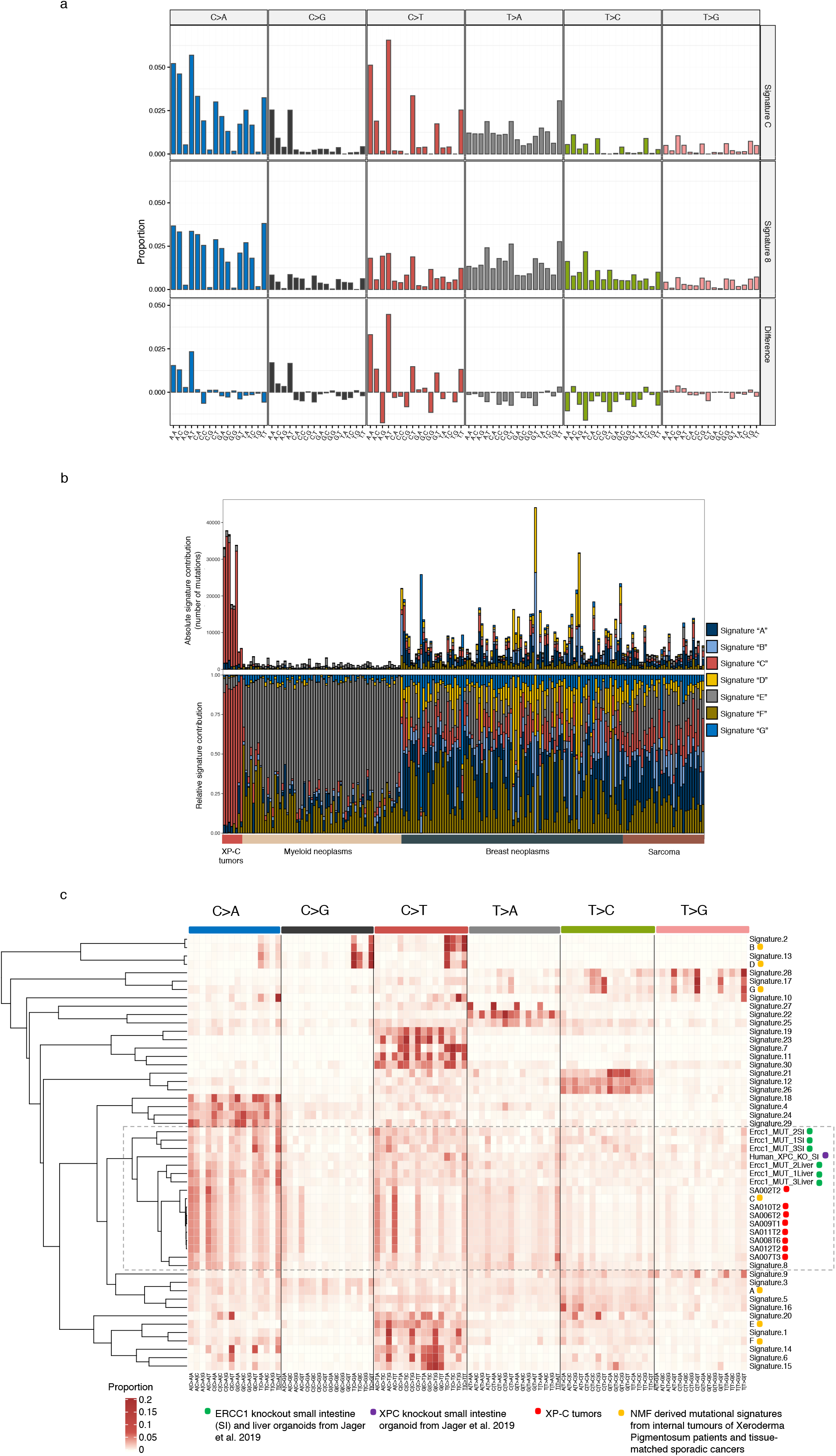
Mutational profiles of XP-C tumors in the context of known mutational signatures. **a**, NMF-derived mutational Signature “C” from XP-C tumors and tissue-matched sporadic cancers in comparison with COSMIC Signature 8 (Alexandrov et al., 2013b) (Cosine similarity = 0.86). **b**, Relative contribution of NMF-derived mutational signatures in XP-C and tissue-matched sporadic cancers (NMF approach). XP-C tumor mutational profiles are dominated by Signature “C”, while sporadic cancers by other signatures with relatively small proportion of Signature “C”. **c**, Unsupervised hierarchical clustering based on the Cosine similarity distances between the XP-C tumors mutational profiles, NMF-derived mutational signatures from XPC tumors and tissue-matched sporadic cancers, COSMIC mutational signatures (Signatures 1-30), and *XPC* and *Ercc1* organoid knockouts (Jager et al., 2019). XP-C tumors cluster with each other and COSMIC Signature 8 forming a larger cluster with *Ercc1* and *XPC* organoid knockouts.

These seven extracted signatures (A to G) together with original XP-C mutational profiles were compared with COSMIC mutational signatures (Alexandrov et al., 2013b) and mutational profiles of organoids from human *XPC* and mouse *Ercc1* knockouts (Jager et al., 2019) using unsupervised clustering. This analysis revealed that the XP-C tumor mutational profiles and their NMF-derived mutational Signature “C” had the highest similarity to the COSMIC Signature 8 (cosine similarity of 0.87 – 0.92, and 0.86 respectively) and formed a cluster together with *XPC* and *Ercc1* organoid knockouts (Fig. 2c., Supplementary Fig. 2e). At the same time the Signature “C” was different from Signature 8 by strong transcriptional asymmetry, increased mutations from C and decreased mutations from T (1.24 and 1.43 fold respectively) specifically in excess of Vp<u>C</u>pT > <u>D</u> and Np<u>C</u>p<u>T</u> > <u>T</u> (where V designates A,C,T and D – A,G,T; Fig. 2a).

A mutational process associated with XPC-deficiency is expected to demonstrate asymmetry between the transcribed and untranscribed strands of a gene (transcriptional bias: TRB) (Zheng et al., 2014). This may be associated with excess of unrepaired bulky lesions on the untranscribed strand due to impaired GG-NER while on the transcribed strand such lesions would be effectively repaired by TC-NER (Haradhvala et al., 2016). Indeed, transcriptional strand bias in XP-C was strong and highly significant for all six classes of nucleotide substitutions grouped by the reference and mutated nucleotide, while in tissue-matched sporadic cancers it was weak or absent (Fig. 3a,b,c,e., Supplementary Fig. 3a,b,c). Moreover, the strongest transcriptional bias was detected in highly expressed genes of XP-C tumors, reaching 7.34-fold, (Wilcoxon signed-rank test, two-sided, *P* = 2.91e-11) in XP-C leukemia (Fig. 3c., Supplementary Fig. 3d).

**Fig. 3.**
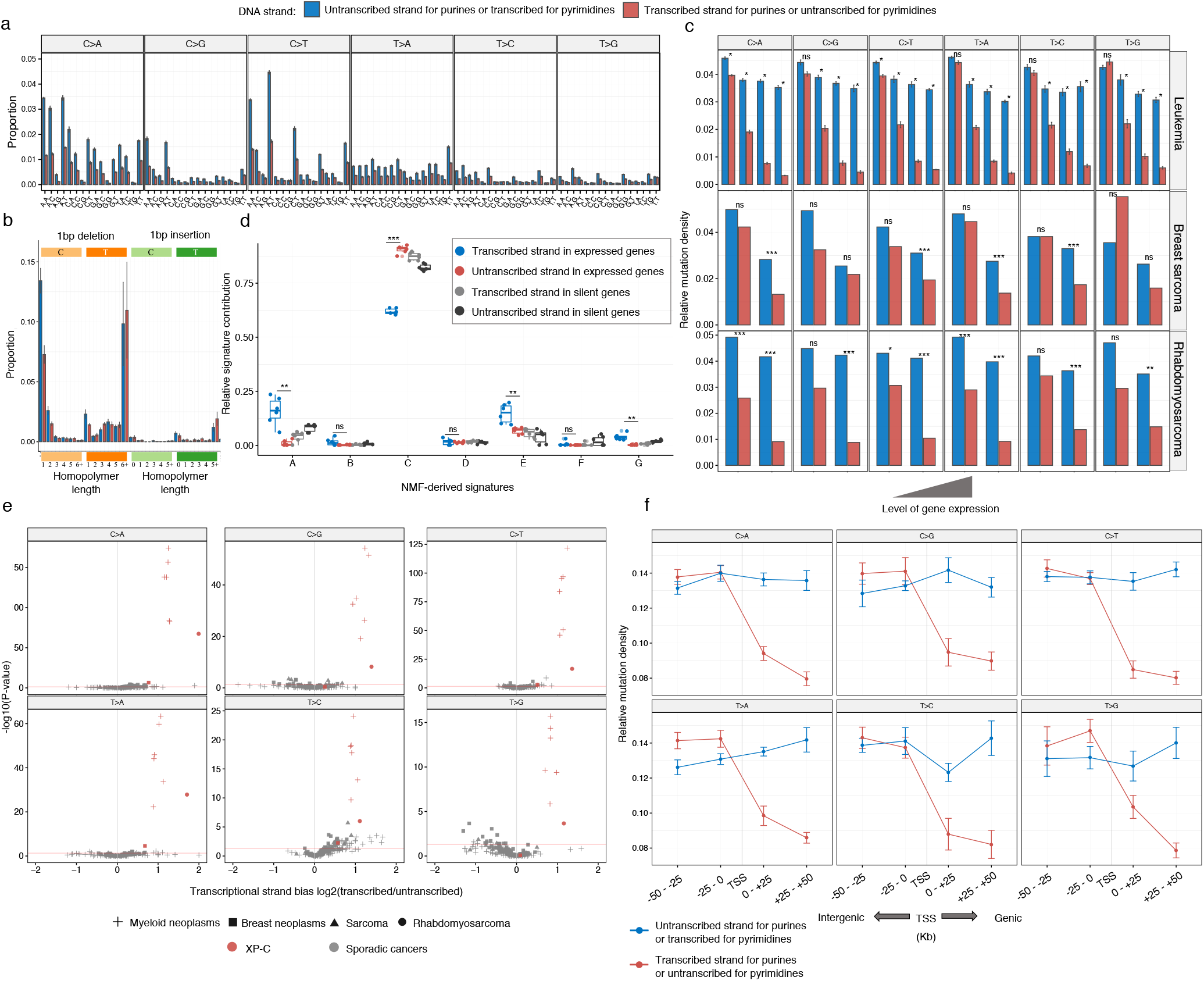
Strong transcriptional bias (TRB) is a specific feature of XP-C tumors. **a**, TRB is observed in the majority of trinucleotide contexts of XP-C leukemia samples (n=6, SEMs are indicated). **b**, TRB is highly pronounced for specific single nucleotide C:G deletions in XP-C leukemia samples (n=6, SEMs are indicated). **c**, TRB strength depends on the level of gene expression and is most pronounced in highly expressed genes (SEMs are indicated for leukemia; Poisson, two-sided test used for breast sarcoma (n=1) and rhabdomyosarcoma (n=1); Wilcoxon signed-rank, two-sided test for leukemia (n=6), *P*: ns – nonsignificant, * < 0.5, ** < 0.01, *** < 0.001). **d,**Relative mutational signature contribution for mutations separated by transcribed and untranscribed strands in transcriptionally active (FPKM > 2) and silent genes (FPKM < 0.05) of XP-C leukemia. Predominant in XP-C leukemia Signature “C” is depleted on the transcribed strands with functional TC-NER, but relative contribution of signatures “A” and “E” typical for sporadic leukemia is enriched on the transcribed strand (T-test, two-sided, paired between transcribed and untranscribed strands in expressed genes (n=6), *P:* ns – nonsignificant, * < 0.5, ** < 0.01, *** < 0.001). **e**, TRB is highly significant and pronounced in XP-C samples for all 6 substitution classes in comparison with sporadic cancers (Poisson two-sided test). **f**, The strong TRB observed in XP-C cancers is caused by transcriptional-coupled repair (TC-NER) but not transcriptional-associated damage. Strong decrease of mutation rate is observed on the genic untranscibed strand for pyrimidines (transcribed for purines, red; right side of transcription start site, TSS), but not on the transcribed strand for pyrimidines (untranscribed for purines, blue) as compared to neighboring intergenic regions (±50 kbp from transcription start site).

These effects could be explained by either excess of mutations from damaged pyrimidines or decrease of mutations from damaged purines on the transcribed (noncoding) strand. Both phenomena were previously described (see Haradhvala et al., 2018) and refer to Transcription-coupled Damage (TCD) or Transcription-coupled repair (TCR). In case of TCD the increase of mutation rates in gene as compared to intergenic region should be observed (TCD in liver cancer analysis in Haradhvala et al., 2018) while in case of TCR we can expect the decrease of mutation rates in gene as compared to intergenic regions. In order to discriminate between these two possibilities a comparison between mutation rates in intergenic and genic regions separately for purines and pyrimidines can be performed. To validate the suspected effect of TC-NER (decrease of mutations from purines on the transcribed strand), we performed two analysis. First, we compared relative signature contributions on the transcribed and untranscribed strands of genes and observed strong depletion of the predominant in XP-C leukemia Signature “C” as well as increase of typical for sporadic leukemia Signatures “A” and “E” on the transcribed strand of genes (Fig. 3d). Second, we compared mutation rates separately on transcribed and untranscribed strands of genes with proximal intergenic regions and observed a strong and significant effect compatible with the decrease of mutations from purines on the transcribed strand (average 1.64 fold, Wilcoxon signed-rank test, two-sided, *P* = 1.694e-13) while there was no significant difference of mutations from purines between intergenic regions and untranscribed strand (*P* = 0.4437; conventional mutation representation depicts decrease of mutations from pyrimidines on the untranscribed strand; Fig. 3f, Supplementary Fig. 3e). In line with that, we observed no difference between mutations from purines on untranscribed strand and intergenic regions at different replication times, while signature of repair of mutations from purines on transcribed strand was observed and was the strongest in early-replicating regions which are usually associated with active gene transcription (Cowie et al., 2014; Zheng et al., 2014) (Fig. 4a, Supplementary Fig. 4a). Similarly to SBS, transcriptional bias in DBS and ID indicated that the primary damage is on purine bases, specifically in <u>CpC</u>>ApD and single nucleotide deletion of C:G nucleotides (Fig. 3b, Supplementary Fig. 3b,c).

**Fig. 4.**
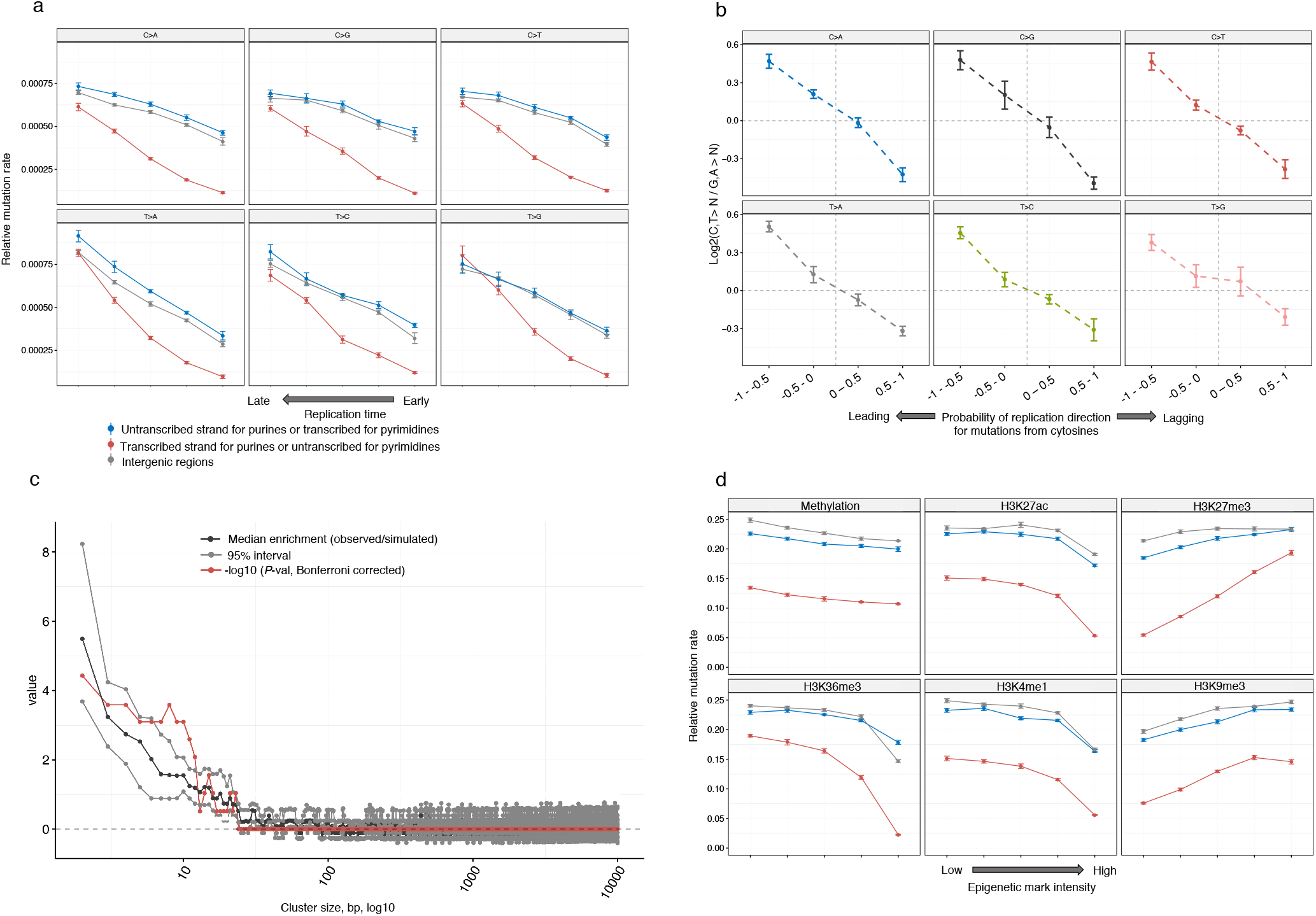
Genomic landscape of mutagenesis in XP-C internal tumors. **a**, Mutational density on the transcribed, untranscribed DNA strands of genes as well in intergenic regions presented as a function of replication timing in XP-C leukemia. Replication time is split onto 5 quantiles. Mutation rate for pyrimidines on the transcribed strand (or purines on the untranscribed, blue) is not different from intergenic regions within the same bin, which is compatible with the absence of GG-NER. **b**, Pyrimidine/purine ratios of mutation rates for regions of the genome grouped by propensity of reference DNA strand to be replicated as leading (left) or lagging (right) strand during DNA synthesis. Strong enrichment of mutagenesis on the leading strand from pyrimidines (C and T) (lagging strands from purines (G and A)) is observed for all 6 classes of mutations. Mutations from purines on the lagging DNA strand may result from error-prone Translesion Synthesis. **c**, The assessment of the length of clustered mutation events on the distances ranging between 2 and 10000bp. Sliding window of 5bp was used to estimate median effect size (black) and its 95% interval (grey) as well as Bonferroni-corrected −log10 (*P*-value) (red, Wilcoxon signed-rank test, two-sided) for different length of clusters in real data (XP-C leukemia, n=6) against simulations (Methods section). The highest and significant enrichment of clustered mutations was observed for short clusters with lengths between 2 and 16 bp. **d**, Intensity of epigenetic marks (5 quantiles) and relative mutation load. Mutational density on the transcribed, untranscribed DNA strands of genes as well in intergenic regions positively correlated with repressive histone marks H3K27me3 and H3K9me3, and inversely correlated with active chromatin marks (H3K27ac, H3K36me3, H3K4me1). For all three genomic categories, effect of the majority of epigenetic marks was similar. At the same time correlations of untranscribed strand for pyrimidines (or transcribed for purines, red) with H3K27me3 and H34M36me3 were more important than for the other two categories.

Recent report suggested that bulky DNA lesions on the lagging strand during DNA replication are more frequently converted into mutations than on the leading strand probably due to more frequent error-prone bypass by translesion synthesis (TLS) polymerases (Haradhvala et al., 2016; Seplyarskiy et al., 2018). Indeed, we found a strong replicational bias (average 1.38-fold of all six mutational classes in XP-C leukemia, Wilcoxon signed-rank test, two-sided, *P* = 2.91e-11) compatible with preferential bypass of purine DNA lesions by error-prone TLS polymerases on the lagging strand (Fig. 4b, Supplementary Fig. 4c) in XPC-deficient tumors.

TLS polymerases which are recruited to bypass a bulky lesion can also insert incorrect bases opposite to undamaged nucleotides near the lesion (Matsuda et al., 2001; Stone et al., 2012). Indeed, in all 8 XP-C tumors we observed statistically significant excess of clustered events as compared to the random distribution (Fig. 4c, Supplementary Fig. 5). In diploid genome regions of XP-C leukemia 0.3% of SBS formed 140 short clusters with distance between mutations inferior to 16bp and mean of 7 bp (Fig. 4c; Supplementary Fig. 5). Moreover, 6.56-fold more mutations, which occurred within a distance of 16bp from each other were co-localized on the same sequencing reads, indicating that clustered mutations affect the same allele and may be interconnected (Wilcoxon signed-rank test, two-sided, *P* = 0.031). These results are compatible with the hypothesis of the existence of bulky DNA lesions which enter the S phase and get bypassed by error-prone translesion DNA synthesis polymerases (Seplyarskiy et al., 2018) in *XPC*-deficient cells, while in *XPC*-proficient cells majority of these lesions may be repaired prior to replication in error-free manner.

Due to the absence of GG-NER we expected to observe strong difference in terms of mutation rates between transcribed and untranscribed strands, particularly in open chromatin and early replicating regions known to be actively transcribed while we expected no difference between untranscribed strand of genes and intergenic regions in heterochromatic regions (Zheng et al. 2014). In XP-C leukemia mutation load in regions of open chromatin was strongly depleted in early replicating regions and regions with active histone marks (H3K27ac (2.83 fold), H3K36me3 (8.45 fold), H3K4me1 (2.72 fold)) for transcribed strands of genes (Figs. 4a,d). Similar but weaker trends were observed when only untranscribed strands of genes and intergenic regions were analyzed (Figs. 4a,d, Supplementary Fig. 4a). Mutation load was also enriched on the untranscribed strand of genes and intergenic regions with repressive histone marks (H3K27me3 (1.26 and 1.09 fold), H3K9me3 (1.28 and 1.25 fold)) and in late replicating regions associated with heterochromatin (Figs. 4a,d). The observed patterns further confirm effectiveness of TC-NER on transcribed strand of genes in euchromatic regions while prove GG-NER being dysfunctional on both intergenic regions and untranscribed strands of genes all over the genome in XP-C samples. To assess the relative mutation rates in different chromatin state regions we compared XP-C leukemia samples and sporadic myeloid neoplasms. The analysis revealed more homogeneous mutation load across the different states in XP-C leukemia in comparison with sporadic leukemia as well as elevated mutation rates in heterochromatic regions relative to genic and regulatory elements (Supplementary Fig. 4b).

To further validate mutational consequences of XPC deficiency, we compared the mutational landscape of cutaneous squamous cell carcinomas (cSCC) from XP-C patients and sporadic tumors (Zheng et al. 2014). All cSCC tumors, independently of XP-C mutational status presented the typical UV-light induced signature (C>T mutations at YpC sites (where Y designates C or T), 85.6%, Supplementary Fig. 6a), which arises due to the bulky lesions on pyrimidines. However, in XP-C cSCCs there was remarkably more pronounced decrease of mutations from pyrimidines on the transcribed strand relative to untrascribed strand and intergenic regions, as well as much stronger transcriptional bias in highly expressed genes (Supplementary Fig. 6b, c). Moreover XP-C cSCC demonstrated stronger difference than sporadic cSCC between mutation rate on the transcribed strand of genes on the one side, and untranscribed strand of genes and intergenic regions on the other (Supplementary Fig. 6b, c, d). These differences were particularly strong in transcriptionally active early replicating regions (Supplementary Fig. 6d). In the case of XP-C internal tumors the observed patterns were similar with the only difference that the mutational profiles are compatible with mutations from purines (Figs. 3c,f; Fig. 4a).

In order to assess the timing of somatic mutations in XP-C tumors we selected the regions of somatic copy number alterations (SCNAs) where one allele was duplicated. We quantified the number of mutations which occurred before and after SCNA (Jolly and Van Loo, 2018) based on variant allele frequencies (n=2307 mutations in 4 copy neutral LOH and 4 copy gains; Supplementary Table 2, Supplementary Fig. 7). On average 75% of mutations occurred before SCNAs suggesting that they may have accumulated in progenitor cells before tumorigenesis or early in tumor development (Wilcoxon signed-rank test, two-sided, *P* = 0.03906; Fig. 5a). Therefore, the observed mutational burden and signature in XP-C tumor genomes may partially represent mutagenesis associated with lesion accumulation during the lifetime of normal body cells (Fig. 5b).

**Fig. 5.**
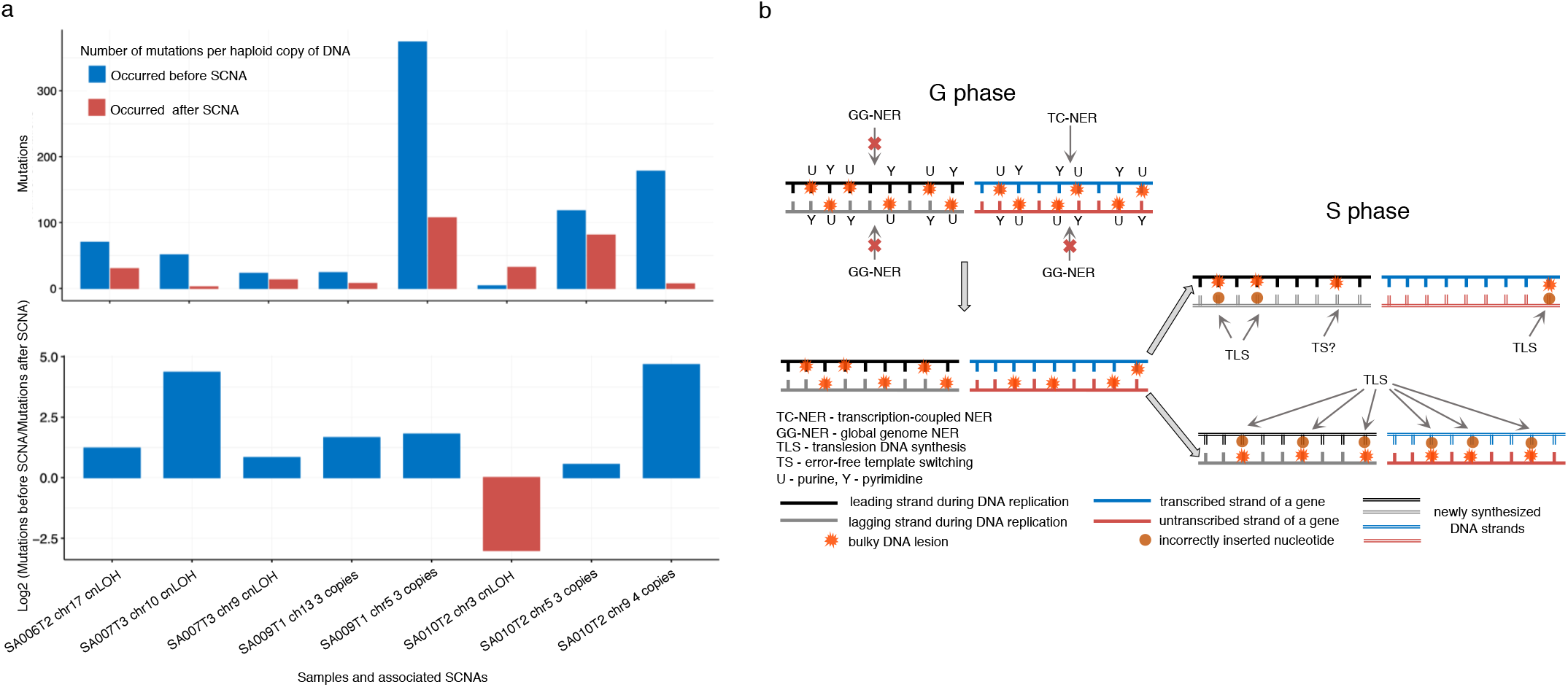
Accumulation of DNA lesions and mutations observed in XP-C tumors. **a**, Relative number of mutations which occurred before and after SCNAs in XP-C cancer genomes (normalized per haploid DNA copy number). The majority of events demonstrate an excess of mutations that were accumulated before the SCNA and may have occurred in tumor-progenitor cells or at early stages of carcinogenesis. **b**, A model of DNA lesion accumulation and mutagenesis in XP-C cells. In XP-C cells where GG-NER is dysfunctional, bulky lesions cannot be efficiently repaired and persist everywhere in the genome except transcribed strands of active genes where TC-NER is operative. During the S-phase a part of bulky lesions on the leading strand may be removed by error-free template switching (TS) mechanisms while on the lagging strand they are converted to mutations by error-prone translesion synthesis (TLS) more frequently, causing mutation accumulation with cell divisions and observed transcriptional and replication biases.

## DISCUSSION

This described mutator phenotype may explain the increased risk of internal cancers in general and particularly for hematological malignancies in XP-C patients, which may be associated with relatively high rate of blood stem cell divisions (Tomasetti, 2016). Our results are in line with recent reports in human and mice showing that attenuated NER at germinal level is associated with increased risk of lymphoma and sarcoma (Chan et al., 2017; Hyka-Nouspikel and Nouspikel, 2011).

The newly derived from XP-C cancers Signature “C” has the highest similarity to COSMIC Signature 8 which was originally extracted from sporadic tumors with the most elevated (but not usually exceeding 35%) fraction in sarcoma, medulloblastoma, lymphoma, chronic lymphocytic leukemia and breast cancer (Alexandrov et al., 2013b). While in some works, it was attributed to homology-repair deficiency (Ma et al., 2018; Waszak et al., 2018), recently in organoid models Signature 8 was associated with the nucleotide-excision repair deficiency (Jager et al., 2019). Comparison of the mutational profiles and NMF-extracted Signature “C” from XP-C internal tumors with the mutational profiles of human *XPC* and mouse *Ercc1* knockouts demonstrated high similarity between them highlighting the dysfunctional NER as the genetic basis of their common mutational process. Our work provides evidence that COSMIC Signature 8 is likely to result from mutagenesis associated with bulky lesions primarily repaired by NER and can be considered as a marker of attenuated NER function.

Taken together our results and previous reports demonstrate that NER-deficiency in different tissue types and in in-vitro models unmasks a unique mutational process of similar etiology. A broad spectrum of nucleotide substitutions and deletions in XP-C context suggests the existence of a compendium of different bulky lesions induced by one or more genotoxins in DNA of somatic cells. The studied patients were diagnosed as XP-C at early age (median: 3 years) and were well protected from environmental mutagens during their life, therefore the observed mutagenesis could be caused by endogenous genotoxins which DNA lesions are almost fully repaired in *XPC*-proficient cells (Fig. 5b).

Future studies on the identification of nature of this mutational process and its link with particular genotoxins (for ex. free radicals, aldehydes, food mutagens) producing bulky lesions may result in elaboration of preventive measures for XP patients. Except the breast sarcoma sample from Comorian Archipelago with IVS12 mutation, our dataset mainly represents a XP-C population of the Northern african origin and single *XPC* mutation (delTG) urging the importance of expanding the investigation of internal tumorigenesis and underlaying mutagenesis in different XP populations.

## ACKNOWLEDGEMENTS

S.N. was supported by grant Foundation ARC 2017, Foundation Gustave Roussy and Swiss Cancer League KFC-3985-08-2016. The authors would like to thank Dr. Patricia Kannouche and Dr. V. B. Seplyarskiy for fruitful discussions and participation, and Dr. F. Rajabi, Dr. Catherine Genestie and Dr. Samuel Quentin for DNA extraction and providing samples. The authors are also very thankful to Dr. C. Genestie (IGR, Villejuif, France), Dr. Z. Tata (Algiers, Algeria) and Dr. S. Duquenne and Dr. F. Cartault (Saint-Pierre, La Réunion, France) for giving us or for manipulating biopsies of tumors and Xiaole Xu (BGI) for perfect management of sequencing.

## AUTHOR CONTRIBUTIONS

S.N., A.S. and A.A.Y. designed the study. A.S. and J.S. collected the samples. A.A.Y performed the data analysis and prepared figures. B.T.M participated in the data analysis. I.P performed data preprocessing. A.A.Y. and S.N. drafted manuscript. A.S. and J.S. commented manuscript. All authors contributed to the final version of the paper.

## SUPPLEMENTARY FIGURES AND TABLES

**Supplementary Table 1. Information about samples used in the study.**

**Supplementary Table 2. Number of mutations occurred before and after SCNAs.**

**Supplementary Table 3. Matrix of single base substitutions (SBS) in different mutational classes.**

**Supplementary Table 4. Matrix of double base substitutions (DBS) in different mutational classes.**

**Supplementary Table 5. Matrix of indels (ID) in different mutational classes.**

**Supplementary Figure 1.**
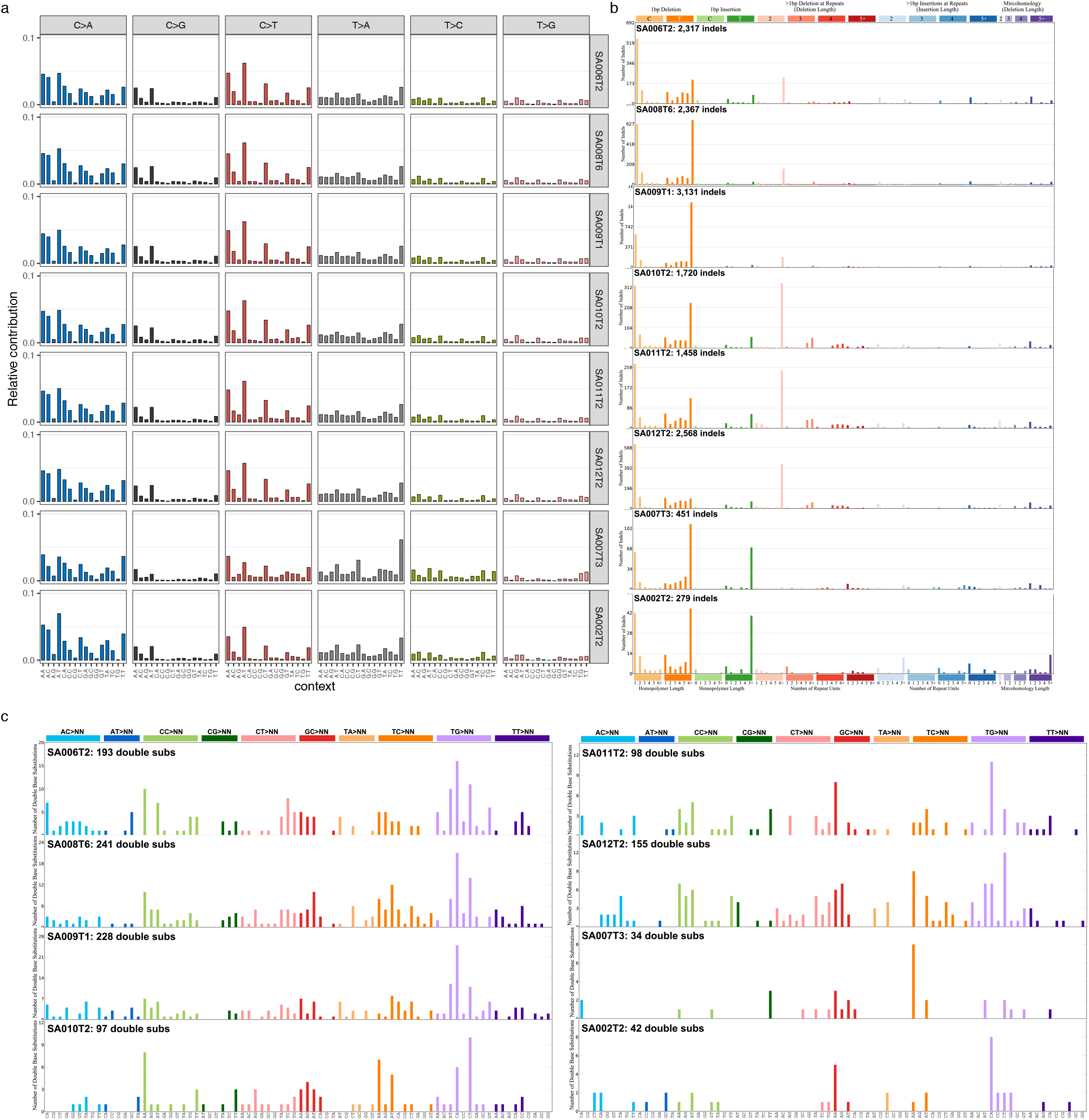
Individual mutational profiles of XP-C tumors for different types of mutations. **a**, Single base substitutions (SBS). **b**, Indels (ID). **c**, Double base substitutions (DBS).

**Supplementary Figure 2.**
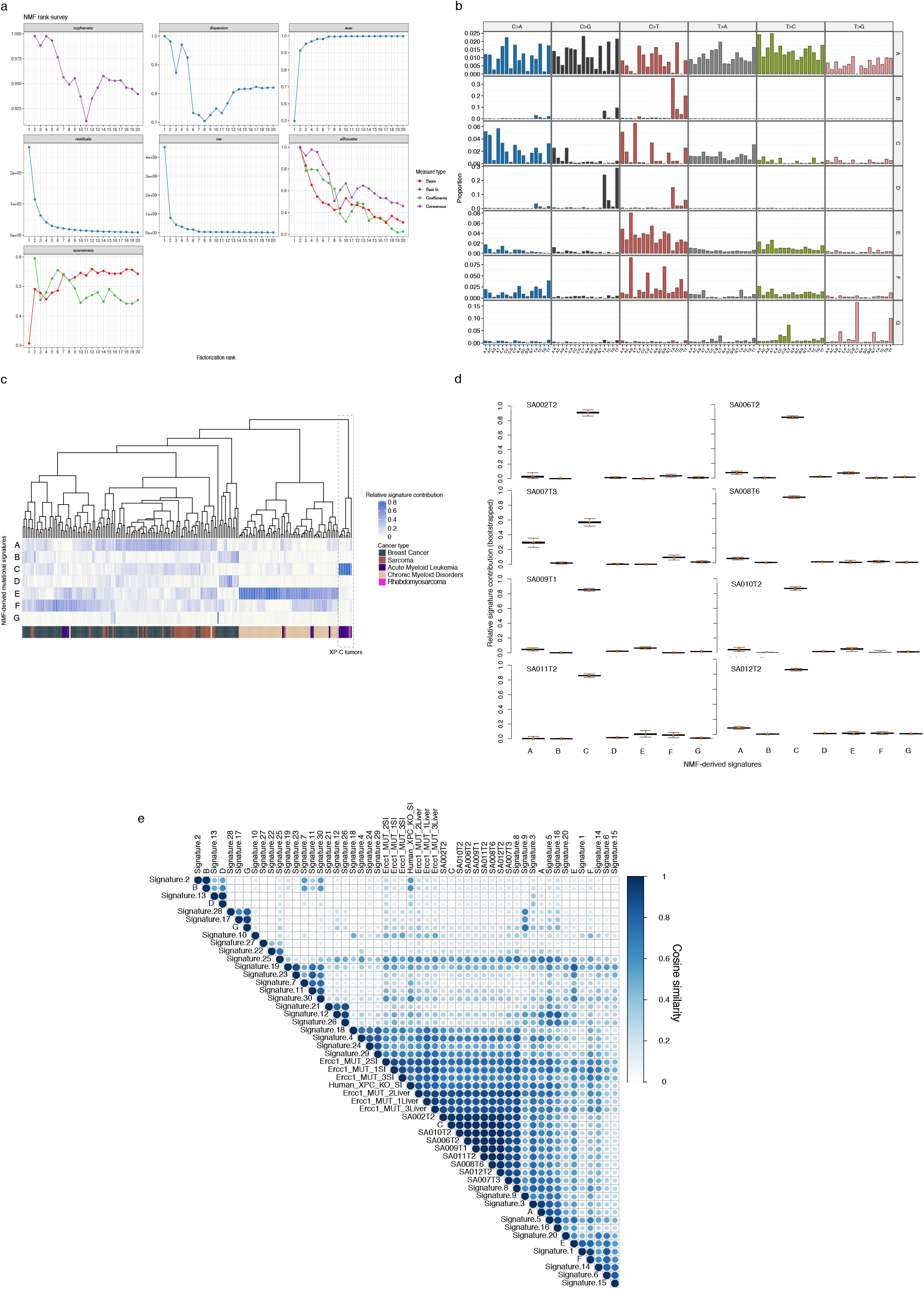
Non-negative Matrix Factorization-derived (NMF) mutational profiles and their contribution in XP-C and sporadic cancers. **a**, Factorization ranks of NMF and diagnostic plots. The model (K=7) was chosen based on the inflation of RSS (Residual Sum of Squares) and evar (explained variance achieved by a model). **b**, Trinucleotide profiles of NMF-derived mutational signatures. **c**, Relative contribution of the NMF-derived mutational signatures in XP-C tumors and sporadic tissue-matched samples (unsupervised hierarchical clustering, relative contribution was inferred using quadratic programming-based algorithm (Huang et al., 2018)). XP-C tumors group together, and their mutational profiles are characterized by high level of Signature “C”. **d**, Bootstrapped estimation (10000 replications) of relative contribution of the NMF-derived signatures in XP-C tumors (quadratic programming-based algorithm). Signature “C” dominates in all the samples with very little level of variation and only in breast sarcoma sample (SA007T3) is slightly depleted. **e**, Cosine similarity matrix between NMF-derived mutational signatures (A-G), COSMIC mutational signatures (Signatures 1-30), mutational profiles of XP-C tumors (SA00..), and *Ercc1* and *XPC* deficient organoid cultures.

**Supplementary Figure 3.**
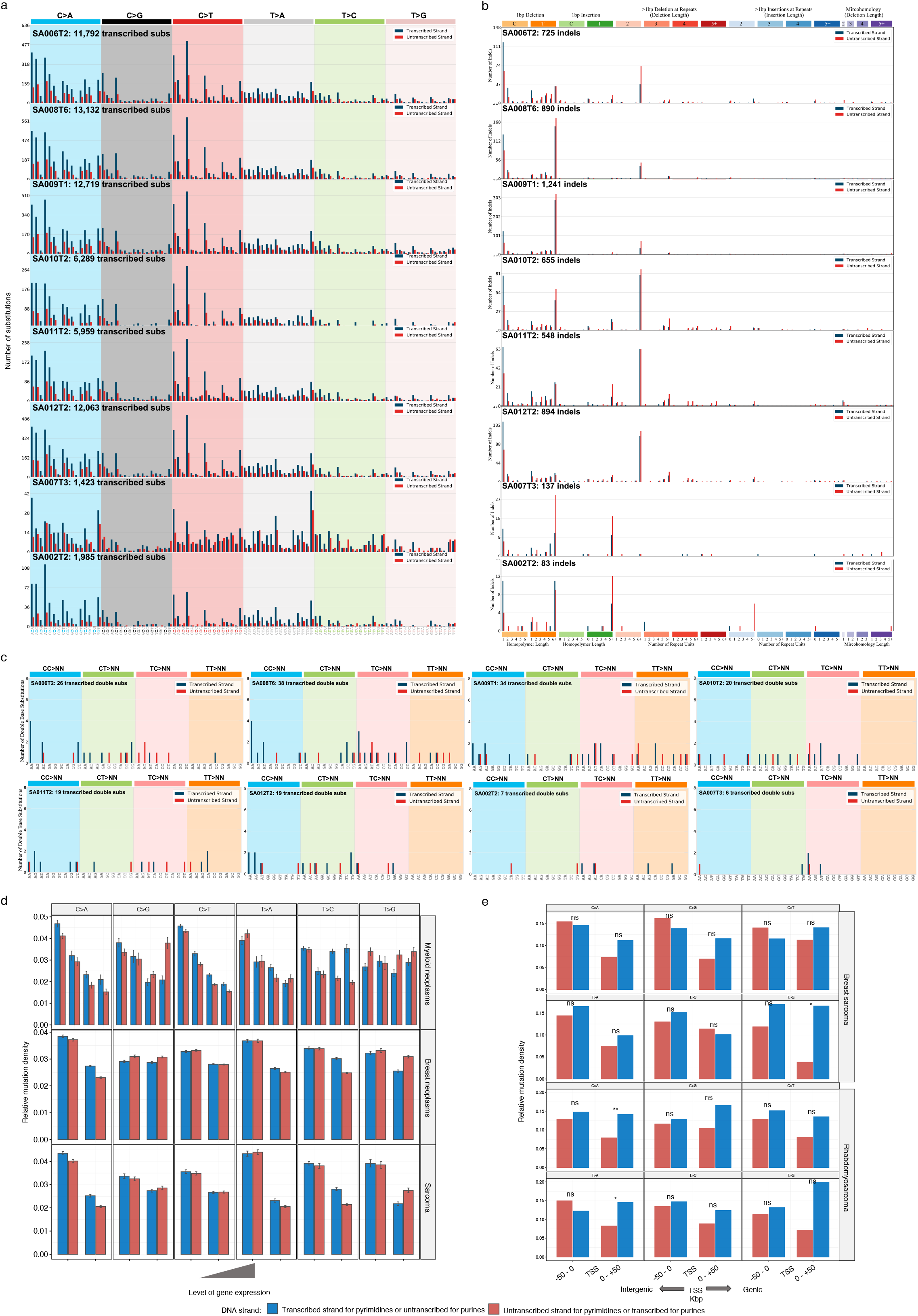
Transcriptional bias (TRB) in XP-C tumors and sporadic cancers. **a,**Stranded mutational profiles of XP-C tumors in genic regions for single base substitutions (SBS). Canonical notation depicts mutations from pyrimidines (blue – transcribed for mutations from pyrimidines, untranscribed for mutations from purines; red – untranscribed for mutations from pyrimidines, transcribed for mutations from purines). **b**, Stranded mutational profiles of XP-C tumors in genic regions for indels (ID). **c**, Stranded mutational profiles of XP-C tumors in genic regions for double base substitutions (DBS). **d**, TRB does not change significantly with the level of gene expression in sporadic tumors (SEM are indicated). **e**, Relative mutation density for mutations from purines and pyrimidines in genic regions and neighboring intergenic regions of XP-C samples (SA007T3 and SA002T2, breast sarcoma and rhabdomyosarcoma respectively, Poisson two-sided test, *P:* ns – nonsignificant, * < 0.5, ** < 0.01, *** < 0.001).

**Supplementary Figure 4.**
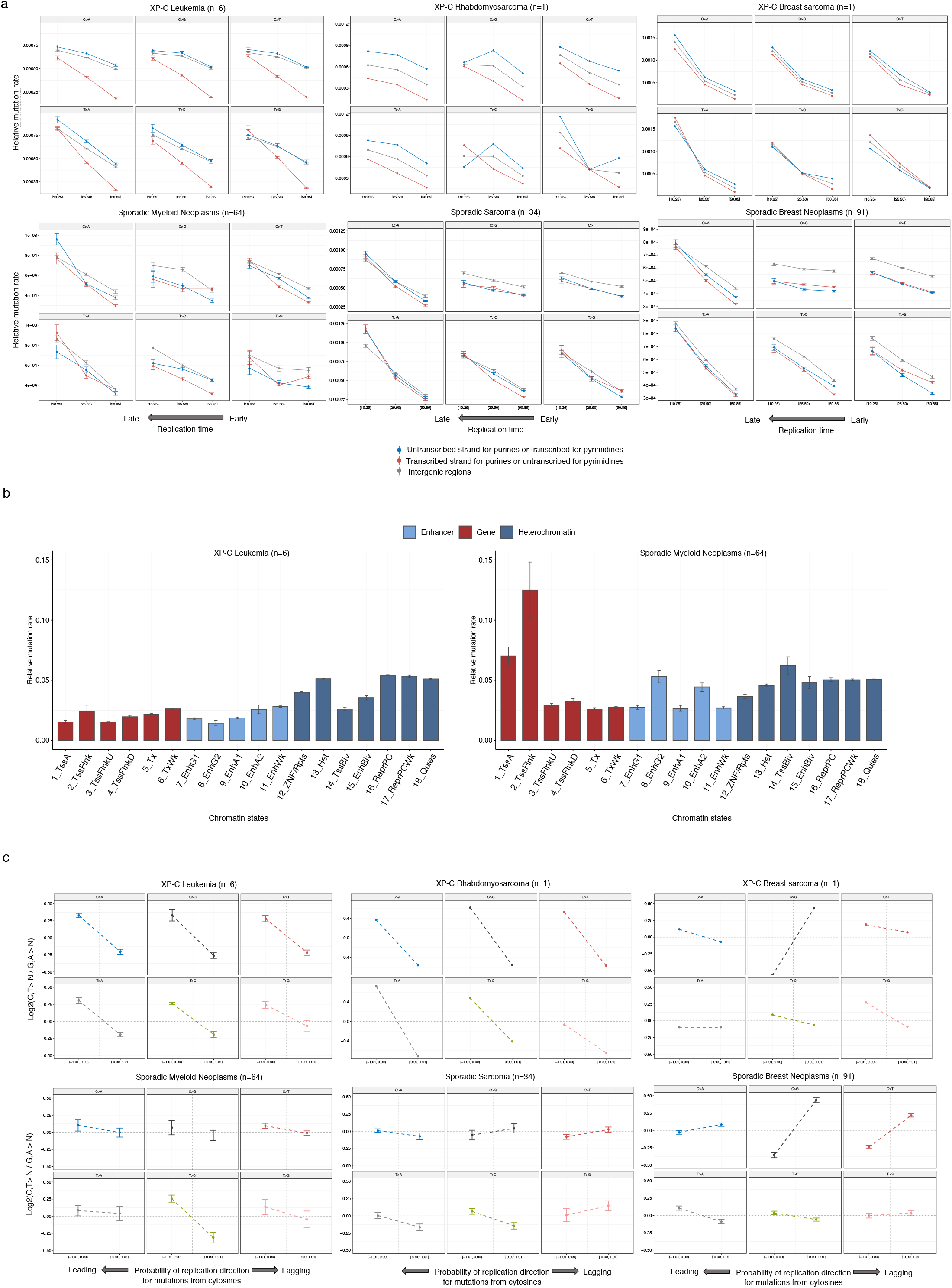
Genomic landscape of mutagenesis in XP-C internal tumors. **a**, Replication timing and intensity of mutagenesis on the transcribed and untranscribed DNA strands as well in intergenic regions of XP-C samples and tissue-matched sporadic cancers. The transcribed strand for pyrimidines (or untranscribed for purines, blue) behaves as intergenic regions genome-wide due to the absence of GG-NER in XP-C tumors (except breast sarcoma, with relatively low amount of Signature “C”) while this effect is very weak in sporadic cancers due to the different mutagenesis process and functional GG-NER. b, Relative mutation rate in different chromatin states (ChromHMM) for XP-C leukemia and sporadic myeloid neoplasms. **c**, Replication direction (leading and lagging) and relative intensity of mutagenesis (replication bias) in XP-C and tissue-matched sporadic tumors. Enrichment of the relative mutagenesis on the leading strand from pyrimidines (C and T) corresponds to the enrichment on the lagging strands from purines (G and A). For all six mutational classes we observe strong enrichment of mutations from purines on the lagging DNA strand of XP-C leukemia and rhabdomyosarcoma samples. The effect is evident but less pronounced in XP-C breast sarcoma. In sporadic cancers the effect is weak or work in opposite direction.

**Supplementary Figure 5.**
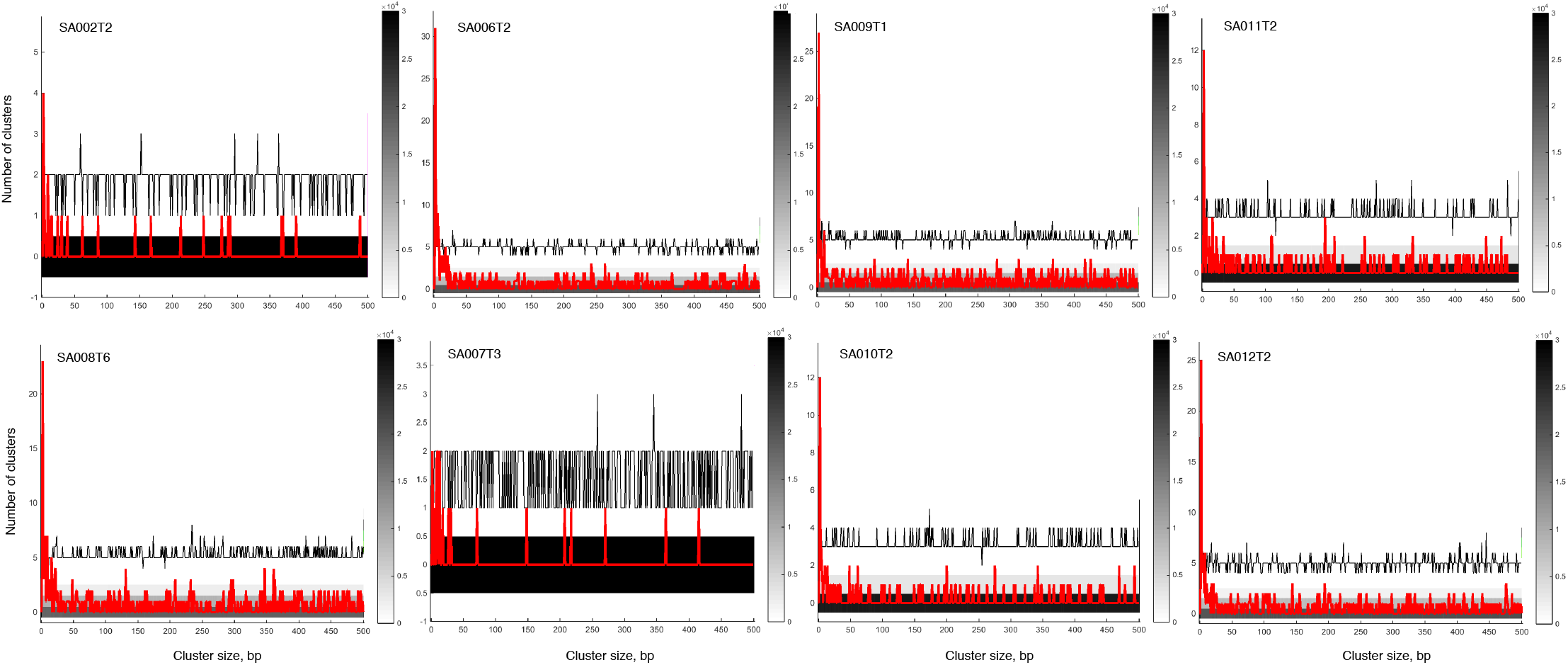
The assessment of the length of clustered mutation events for 1-500bp distances. Observed inter-mutation distance (red) is compared to the density distribution of 30000 simulations (black) with similar number of mutations and trinucleotide contexts. Strong enrichment of clustered events is evident at short cluster distances in the all the samples.

**Supplementary Figure 6.**
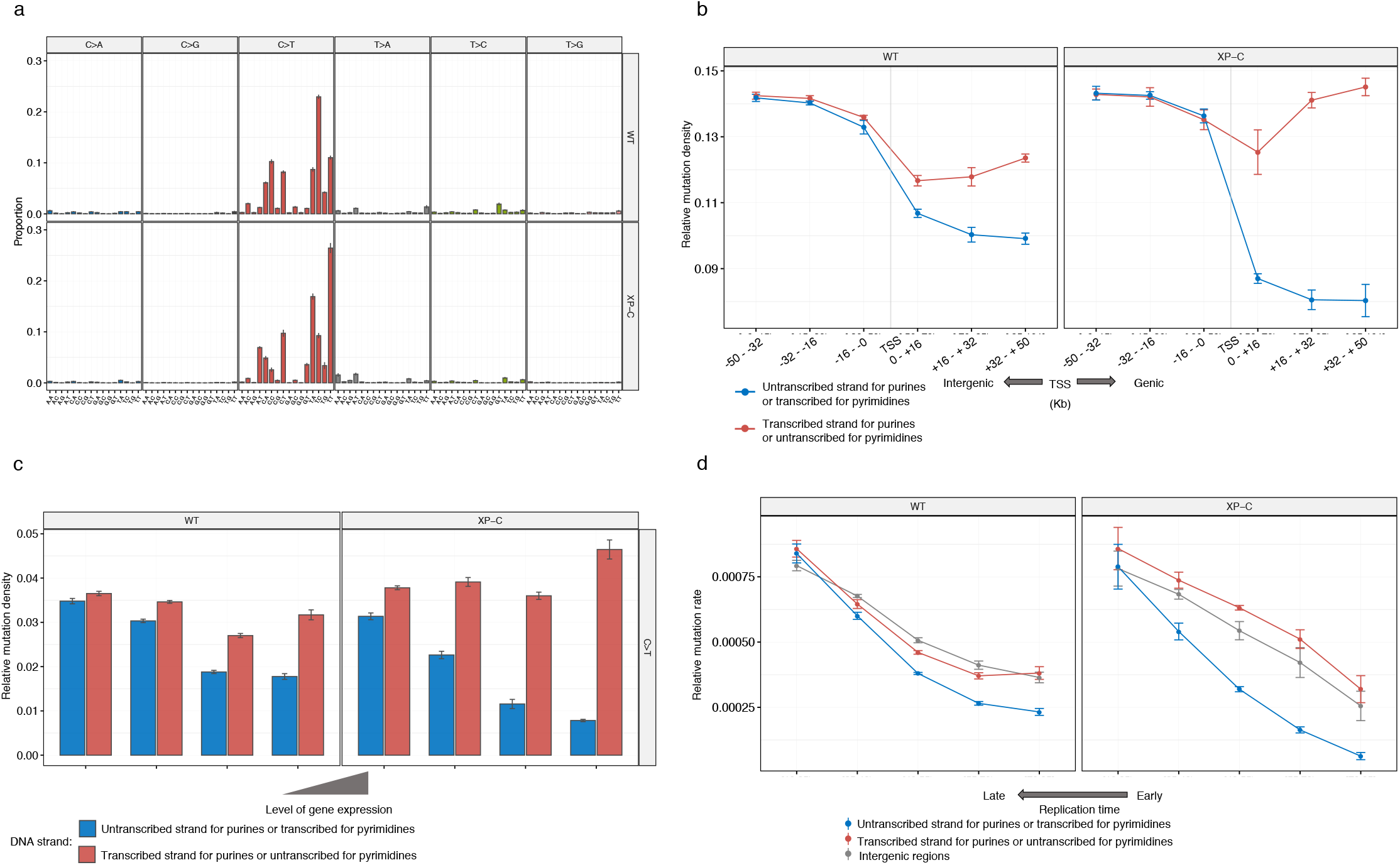
Genomic mutational landscape of WT (n=7) and XP-C (n=5) cutaneous squamous cell carcinoma (cSCC) samples. **a**, Trinucleotide-context mutational profiles (SEM intervals are shown). X-axis represents the nucleotides upstream and downstream of mutation. The mutational profiles of both WT and XP-C cSCCs are dominated by C>T mutations at YpC sites (where Y designates C or T). **b**, Relative mutation density for mutations from purines and pyrimidines in genic regions and neighboring intergenic regions of WT and XP-C cSCC sampels. **c**, TRB strength depends on the level of gene expression and is most pronounced in highly expressed genes (SEMs are indicated) specifically in XP-C cSCC samples. **d**, Replication timing and intensity of mutagenesis on the transcribed and untranscribed DNA strands as well in intergenic regions of WT and XP-C cSCC samples. The untranscribed strand for pyrimidines (or transcribed for purines, red) behaves similar to intergenic regions genome-wide due to the absence of GG-NER in XP-C tumors while this effect is weaker in WT samples due to the different mutagenesis process and functional GG-NER.

**Supplementary Figure 7.**
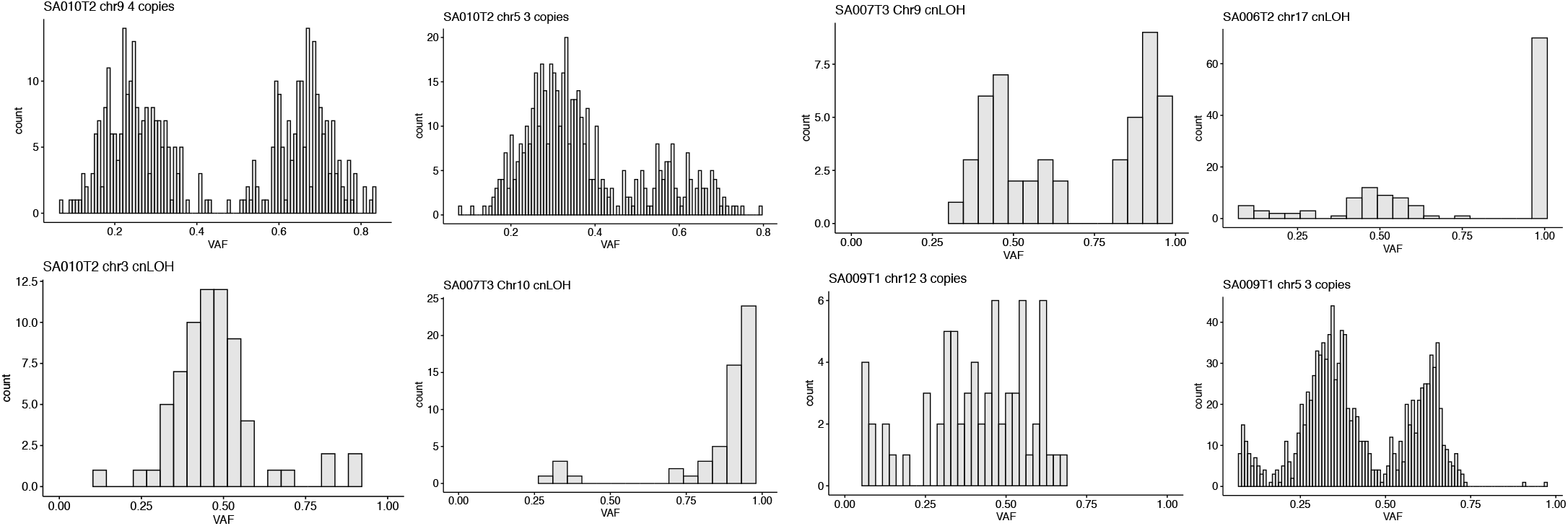
Variant allele frequency distribution in SCNA regions of XP-C tumors.

## METHODS

### Studied samples

Patients from the study were diagnosed with Xeroderma Pigmentosum at early age (median: 3.5 years (range 1.5-9 years); Table 1 and Supplementary Table 1). Primary fibroblasts from sun-unexposed skin were used to determine the DNA repair deficiency by unscheduled DNA synthesis following UV-C irradiation as described (Sarasin et al., 1992). The XP genetic defect was characterized by complementation assay using recombinant retroviruses expressing wild type DNA repair genes (Arnaudeau-Bégard et al., 2003). The absence of the XPC protein was shown by Western blots (Cartault et al., 2011). The *XPC* mutation was determined by Sanger sequencing or whole exome sequencing. Informed signed consents were obtained from patients and/or their parents in accordance with the Declaration of Helsinki and the French law. This study was approved by the French Agency of Biomedicine (Paris, France), by the Ethics Committee from the CPP of Universitary Bordeaux Hospital (Bordeaux, France) and by the Institutional Review Board of the University Institute of Hematology (IUH: Saint-Louis Hospital, Paris). For patients with leukemia (n=6), tumoral bone marrow or peripheral blood mononucleated cells were separated on Fycoll-Hypaque. Cultured skin fibroblast cells were used as non-hematopoietic DNA controls in 5 out of 6 patients. In the additional patient, bone marrow CD34+, CD14+ and CD3+ cells were sorted with magnetic beads; CD34+ CD14+ cells represented the leukemic fraction while CD3+ T lymphocytes, non-leukemic fraction was used as a control. DNA from solid tumors (SA002T2 and SA007T3) was extracted from FFPE blocks after examination and dissection by a pathologist. Tumor DNA was extracted from parts of FFPE containing more than 90% of tumor cells. Germline DNA was extracted from the non-tumoral part of FFPE (Supplementary Table 1).

### Genome sequencing and data processing

The genomes were sequenced using BGISEQ-500 or Illumina Hiseq 2500 (SA008T6) sequencers according to the manufacturer protocols to the mean coverage after deduplication equal to 45x for tumor and 30x for normal DNA (Supplementary Table 1) using 100bp paired-end reads. Reads were mapped using BWA-MEM (v0.7.12) software (Li and Durbin, 2009) to the GRCh37 human reference genome and then used the standard GATK best practice pipeline (Van der Auwera et al., 2002) to process the samples and call somatic genetic variants. PCR duplicates were removed and base quality score recalibrated using GATK (Depristo et al., 2011) (v4.0.10.1), MarkDuplicates and BaseRecalibrator tools. Somatic SNVs and INDELs were called and filtered using GATK tools Mutect2, FilterMutectCalls and FilterByOrientationBias and annotated with oncotator (Ramos et al., 2015) (v1.9.9.0). SCNAs calling was done with FACETS (Shen and Seshan, 2016) (v 0.5.14). Quality controls of fastq and mapping were done with FASTQC (Andrews, 2015) (v0.11.7), samtools (Li et al., 2009) (v1.9), GATK HSmetrics, mosdepth (Pedersen and Quinlan, 2018) (v0.2.5) and multiqc (Ewels et al., 2016) (v1.5). All processing steps were combined in a pipeline built with snakemake (Köster and Rahmann, 2012) (v5.4.0).

The cutaneous squamous cell carcinoma samples (cSCC) from the work of Zheng et al. 2014 were downloaded as SRA files from the database of Genotypes and Phenotypes (dbGaP). The dataset was processed and filtered in the same way as XP-C leukemia samples.

### Filtration of somatic variants

For XP-C leukemia samples from bone marrow biopsies we used additional filtration of the PASS variants which included requirement of at least 1 read on the both strands (F1R2.split (‘,’).1 > 0 && F2R1.split (‘,’).1 > 0 filters in GATK) and the variant allele frequency (VAF) minimal threshold equal to 0.05.

To avoid contamination of true variants by FFPE sequencing artefacts we used more stringent criteria for breast sarcoma (SA007T3) and rhabdomyosarcoma (SA002T2) samples which included at least 2 and 1 reads from each strand and minimal VAF equal to 0.3 and 0.4 for breast cancer and rhabdomyosarcoma samples respectively. These thresholds were chosen empirically taking into account the high purity/ploidy of the samples (Supplementary Table 1) and VAF of FFPE artefacts which can vary between 0.01 and 0.15 (Robbe et al., 2018).

Additionally, all used VCF files were filtered based on the alignability map of human genome (Derrien et al., 2012) from UCSC browser (Kent et al., 2002) (https://genome.ucsc.edu/cgi-bin/hgFileUi?db=hg19&g=wgEncodeMapability) with the length of K-mer equal to 75bp (wgEncodeCrgMapabilityAlign75mer, mutations overlapped regions with score < 1 were filtered out) and UCSC Browser blacklisted regions (Duke and DAC).

### Mutational signatures analysis

To convert the VCF files into a catalog of mutational matrixes we used the MutationalPatterns software v.1.11.0 (Blokzijl et al., 2018). Profiling of the mutational matrixes of indels and double nucleotides substitutions was performed with SigProfilerMatrixGenerator v.1.0 software (Bergstrom et al., 2019).

For comparison with XP-C tumors we used 190 tissue-matched whole cancer genomes from the ICGC PCAWG collection (Yung et al., 2017) which included cancers from the following projects: Chronic Myeloid Disorders – UK (n=57), Acute Myeloid Leukaemia – KR (n=8), Breast Cancer TCGA US (n=91), Sarcoma - TCGA US (n=34). We used only high-quality variants and additionally filtered out mutations in low-mappability and blacklisted regions of the human genome.

To construct the multidimension scaling plot (MDS) we computed pairwise Cosine similarity distance between all pairs of the samples using MutationalPatterns package (Blokzijl et al., 2018) and then processed the matrix of distances between the samples in prcomp () function in R.

To perform Non-negative Matrix Factorization approach and extract de novo mutational signatures we used the XP-C samples along with tissue-matched dataset of PCAWG samples (n = 190) in NMF framework realized in MutatioanlPatterns R package (Blokzijl et al., 2018) with 500 initialization runs. After examination of the diagnostic plots (Supplementary Fig. 2a) we choose K=7 (with RSS at inflation point, according to Hatchins et al. (Hutchins et al., 2008)) to extract mutational signatures (Supplementary Figs. 2b) and then assigned them to the known mutational signatures based on the Cosine similarity (Fig. 2c, Supplementary Fig. 2e). Choosing of lower (K=4) or higher factorization rank (K=9) did not influence significantly the extracted Signature “C” and its proportion in samples (data not shown).

To quantify the contribution of the NMF-derived mutational signatures (A-G) in XP-C tumors and tissue-matched PCAWG cancers we used the quadratic programming-based algorithm (Huang et al., 2018) realized in SigsPack R package (Schumann et al., 2019) (Fig. 2b). To better understand and quantify the contribution of the NMF-derived mutational signatures in XP-C dataset we additionally used bootstrapping (n=10000) on substitution classes to receive the confidence intervals of each signature contribution (Supplementary Fig. 2d)

### Transcriptional strand bias analysis

Transcriptional strand bias (TRB) was quantified for each sample and six mutational classes using MutationalPatterns package (Blokzijl et al., 2018). The function computed inequality between mutations from pyrimidines (C>A,T,G; T>A,C,G) to mutations from purines (G>A,C,T; A>C,G,T) for genes located on the sense and antisense strands of DNA relative to the reference human genome. Inequality in number of mutations from purines and pyrimidines was considered as evidence of transcriptional bias and statistical significance was assessed using Poisson test.

To compute tissue-specific TRB between genes expressed at low and high level we used RPKM values of RNA-seq from Epigenetic Roadmap Project (Consortium et al., 2015) (E028 for breast sarcoma, E050 for leukemia, E100 for rhabdomyosarcoma). For each gene mutations were separated as located on transcribed or untranscribed strands and genes were divided into bins by the level of expression (RPKMs: 0-0.1,0.1-1,1-10,10-20000 for leukemia; 0-0.1,0.1-20000 for breast sarcoma and rhabdomyosarcoma). The significance for each bin was assessed using Poisson test, two-sided (single samples of breast sarcoma and rhabdomyosarcoma) or Wilcoxon signed-rank test, two-sided (leukemia, n=6) and then for visualization the number of mutations were normalized by the total length of genes in each bin.

Following the hypothesis that majority of mutations were caused by purine DNA lesions we were able to compute strand-specific mutation densities around transcription start sites (TSSs). Transcribed and untranscribed strands of genes as well as 5’ adjacent to TSS intergenic regions were treated separately. TSSs of all annotated genes (GENECODE v30 (Frankish et al., 2019)) were retrieved using BEDTools v2.29.0 (Quinlan, 2014) and then regions located ± 50kb of TSSs were split into 1kb intervals. The 1kb intervals which overlapped with other intergenic or genic intervals (represented mainly by overlapped or closely located genes) were removed. This approach rendered 237 Mbp of 5’ proximal to TSS intergenic regions and 151 Mbp of genic regions.

### Replication timing

We used repliseq data from 12 cell lines (Hansen et al., 2010; Thurman et al., 2007) to calculate consensus replication timing regions. For each 1kb regions we calculated standard deviation between all the cell lines and removed all regions with standard deviation higher than 15. For the rest of consistent regions across different cell lines we calculated mean values and used them during analysis. The genome was divided into 5 bins (10-25,25-40,40-55,55-70,70-85) according to the replication timing values and mutational density was calculated for each bin adjusting for the length of each region. We computed dependence of mutational density on replication timing independently for genic and intergenic regions separating mutations on transcribed strand and untranscribed strands.

### Epigenetic marks and mutational density

To infer relationship between mutation density and intensity of various epigenetic marks (methylation, H3K27ac, H3K27me3, H3K36me3, H3K4me1, H3K9me3) we downloaded bigwig files of the Roadmap Epigenomics Project (Consortium et al., 2015) and converted them to wig and then bed files (tissue E050). The mean intensity of each mark was calculated for 1kb nonoverlapping windows across autosomes with BEDOPS v2.4.37 (bedmap) software (Neph et al., 2012). We used only genomic windows with high alignability (equal to 1) along at least 90% of a window. Mark intensities were normalized to 1-100 range. For each window we split mark intensities into 5 quantiles (cut2() function in R (R Development Core Team, 2011)) and calculated relative mutation density of each mark for intergenic regions, transcribed and untranscibed strands of genes.

The ChromHMM Expanded 18-state models of chromatin states (E050) were downloaded as bed file (Consortium et al., 2015) and all the windows with the highest alignability spanning less than 90% of the window were filtered out. Then we calculated relative mutation density for each sample and chromatin state for XP-C leukemia and sporadic myeloid neoplasms.

### Replicational strand bias

We used data from Okazaki-seq experiments data (Petryk et al., 2016) for GM06990 and HeLa cell lines to infer the regions of genome preferentially replicating as lagging or leading strand relative to the reference human genome. 1kbp genomic regions for which values representing direction of replication fork differed between cell lines more than 0.4 were removed. We calculated ratio of the densities between mutations from pyrimidines (C, T) and purines (G, A) for each bin (−1 - 0.5,-0.5 - 0,0 - 0.5, 0.5 - 1) of the preferential replication direction (negative values correspond to genomic regions where reference strand is replicated as lagging strand; and positive values – as leading) similar to the methodology of Seplyarsky et al. (Seplyarskiy et al., 2016)

### Clustered mutations

To evaluate the distribution of mutations across the genome for the presence of clustered mutations in our dataset, we performed Monte Carlo simulations for the intermutation distances distribution of random mutations for ranges between 2 and 10000bp for each studied sample. We developed a mathematical model of the Monte Carlo method for random mutations generation based on the following statements: 1) positions of mutations are random and uniformly distributed along the genome; 2) random positions are selected from the same set of genomic intervals as original somatic mutations; 3) the number and nucleotide context spectrum of randomly generated mutations exactly matches somatic mutations in the corresponding sample. As follows, our simulations are based on the discrete homogeneous Poisson point process. The Monte Carlo simulations were performed using Java programming language, discrete random positions were generated with standard Java Random class (Supplementary code). Data analysis was carried out with MathWorks MATLAB. We randomly assigned mutations giving their trinucleotide (3bp) contexts and repeated the procedure 30000 times for each sample (Supplementary Fig. 5).

To compute statistics for the distances between neighbors for randomly placed mutations within mapability sections for chromosomes and whole genome we used the following algorithm:

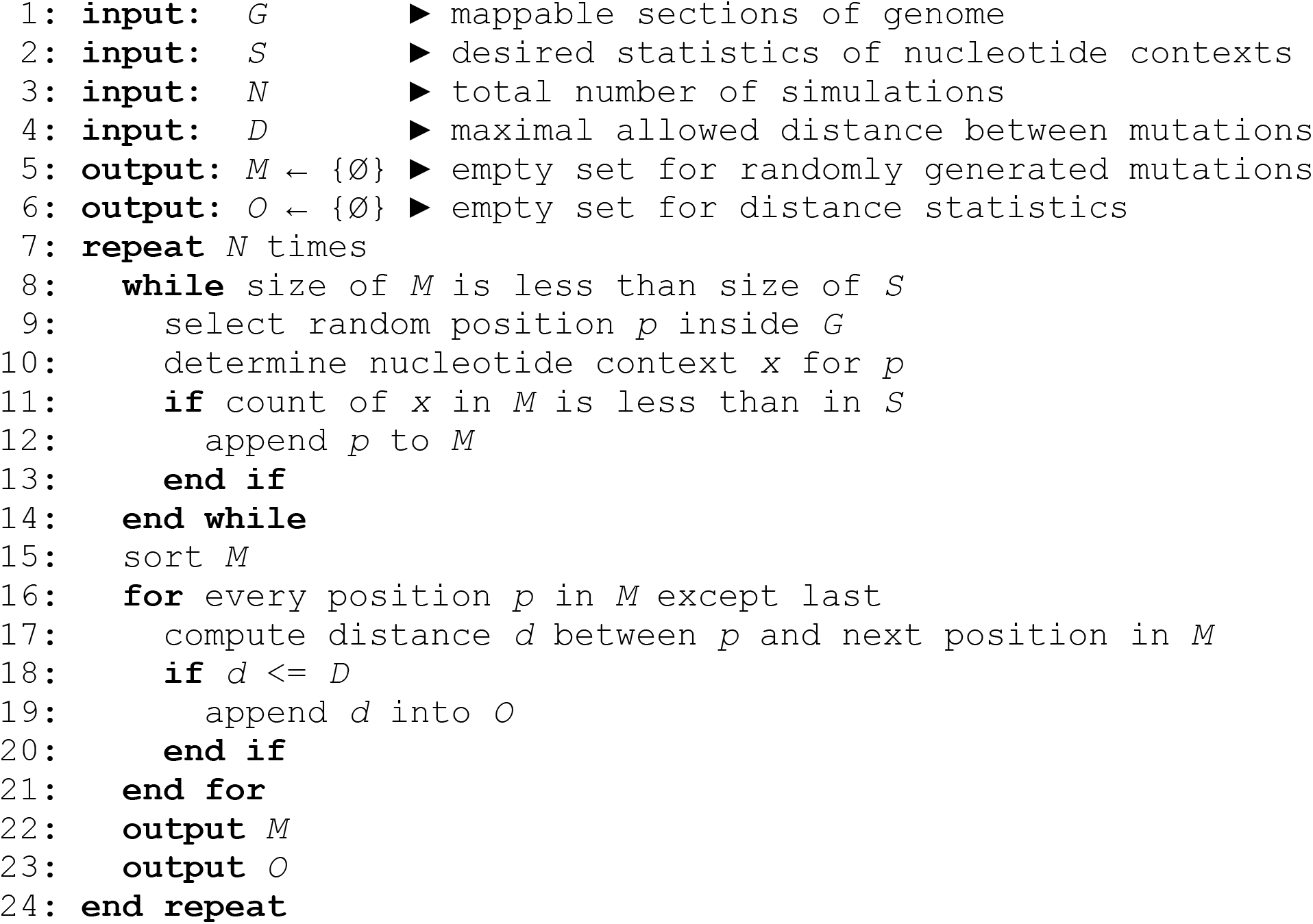

We next verified that random mutations at small distances produced by random generations followed the Poisson distribution. Then, the means for simulated distributions were compared with the observed intermutation distances for XP-C leukemia samples (n=6) using Wilcoxon signed-rank, two-sided test in 5bp overlapping (1bp step) windows to define length of clusters (for 2-10000bp intervals). Resulted P-values were corrected with Bonferroni approach. Significant enrichment of clustered mutations at short distances remained when simulations were performed without taking into account the context of mutations or in 5-bp context of mutations; or when only euploid parts of the genomes were taken into account (data not shown). 4 exomes of XP-C samples were independently sequenced on Illumina Hiseq 2500 with ~100X sequence coverage. Out of 6 clusters that overlapped exonic regions all 6 were validated. Additionally, we assessed the number of mutations located on the same read or different reads for clusters up to 16 bp located in diploid genomic regions.

### Relative number of mutations before and after SCNAs

To infer relative number of mutations which occurred before and after SCNA we followed to the previously described methodology (Jolly and Van Loo, 2018) and identified SCNA of two classes in our dataset: copy gain or cnLOH (Supplementary Table 2). In these SCNA regions taking into account tumor purity and ploidy of the regions we determined conservative variant allele frequency (VAF) thresholds to separate mutations on occurred before and after SCNA given their VAF. The number of mutations was then normalized per haploid copy of a genomic segment.

## DATA AVAILABILITY

Experimental data generated in this study have been deposited to the European Genome-phenome Archive (EGA) accession XXX.

## CODE AVAILABILITY

All software used is published and/or in the public domain. Custom Java code for the clustered mutation analysis is available as the Supplementary code.

## REFERENCES

Alexandrov, L.B., Nik-Zainal, S., Wedge, D.C., Campbell, P.J., and Stratton, M.R. (2013a). Deciphering Signatures of Mutational Processes Operative in Human Cancer. Cell Rep. 3, 246–259.

Alexandrov, L.B., Nik-Zainal, S., Wedge, D.C., Aparicio, S.A.J.R., Behjati, S., Biankin, A. V., Bignell, G.R., Bolli, N., Borg, A., Børresen-Dale, A.L., et al. (2013b). Signatures of mutational processes in human cancer. Nature 500, 415–421.

Andrews, S. (2015). FASTQC A Quality Control tool for High Throughput Sequence Data. Babraham Inst.

Arnaudeau-Bégard, C., Brellier, F., Chevallier-Lagente, O., Hoeijmakers, J., Bernerd, F., Sarasin, A., and Magnaldo, T. (2003). Genetic correction of DNA repair-deficient/cancer-prone xeroderma pigmentosum group C keratinocytes. Hum. Gene Ther.

Van der Auwera, G.A., Carneiro, M.O., Hartl, C., Poplin, R., Del Angel, G., Levy-Moonshine, A., Jordan, T., Shakir, K., Roazen, D., Thibault, J., et al. (2002). GATK Best Practices. Curr. Protoc. Bioinformatics.

Bergstrom, E.N., Huang, M.N., Mahto, U., Barnes, M., Stratton, M.R., Rozen, S.G., and Alexandrov, L.B. (2019). SigProfilerMatrixGenerator: a tool for visualizing and exploring patterns of small mutational events. BMC Genomics 20, 1–12.

Blokzijl, F., Janssen, R., van Boxtel, R., and Cuppen, E. (2018). MutationalPatterns: Comprehensive genome-wide analysis of mutational processes. Genome Med.

Bradford, P.T., Goldstein, A.M., Tamura, D., Khan, S.G., Ueda, T., Boyle, J., Oh, K.S., Imoto, K., Inui, H., Moriwaki, S.I., et al. (2011). Cancer and neurologic degeneration in xeroderma pigmentosum: Long term follow-up characterises the role of DNA repair. J. Med. Genet.

Cartault, F., Nava, C., Malbrunot, A.C., Munier, P., Hebert, J.C., N’guyen, P., Djeridi, N., Pariaud, P., Pariaud, J., Dupuy, A., et al. (2011). A new XPC gene splicing mutation has lead to the highest worldwide prevalence of xeroderma pigmentosum in black Mahori patients. DNA Repair (Amst).

Chan, S.H., Lim, W.K., Ishak, N.D.B., Li, S.T., Goh, W.L., Tan, G.S., Lim, K.H., Teo, M., Young, C.N.C., Malik, S., et al. (2017). Germline Mutations in Cancer Predisposition Genes are Frequent in Sporadic Sarcomas. Sci. Rep. 7, 1–8.

Consortium, R.E., Kundaje, A., Meuleman, W., Ernst, J., Bilenky, M., Yen, A., Heravi-Moussavi, A., Kheradpour, P., Zhang, Z., Wang, J., et al. (2015). Integrative analysis of 111 reference human epigenomes. Nature 518, 317–330.

Cowie, D.A., Nazarethi, J., and Story, D.A. (2014). Chromatin organization is a major influence on regional mutation rates in human cancer cells. Anaesth. Intensive Care 42, 310–314.

Depristo, M.A., Banks, E., Poplin, R., Garimella, K. V., Maguire, J.R., Hartl, C., Philippakis, A.A., Del Angel, G., Rivas, M.A., Hanna, M., et al. (2011). A framework for variation discovery and genotyping using next-generation DNA sequencing data. Nat. Genet.

Derrien, T., Estellé, J., Sola, S.M., Knowles, D.G., Raineri, E., Guigó, R., and Ribeca, P. (2012). Fast computation and applications of genome mappability. PLoS One.

Ewels, P., Magnusson, M., Lundin, S., and Käller, M. (2016). MultiQC: Summarize analysis results for multiple tools and samples in a single report. Bioinformatics.

Frankish, A., Diekhans, M., Ferreira, A.M., Johnson, R., Jungreis, I., Loveland, J., Mudge, J.M., Sisu, C., Wright, J., Armstrong, J., et al. (2019). GENCODE reference annotation for the human and mouse genomes. Nucleic Acids Res.

Hadj-Rabia, S., Oriot, D., Soufir, N., Dufresne, H., Bourrat, E., Mallet, S., Poulhalon, N., Ezzedine, E., Grandchamp, B., Taïeb, A., et al. (2013). Unexpected extradermatological findings in 31 patients with xeroderma pigmentosum type C. Br. J. Dermatol. 168, 1109–1113.

Hansen, R.S., Thomas, S., Sandstrom, R., Canfield, T.K., Thurman, R.E., Weaver, M., Dorschner, M.O., Gartler, S.M., and Stamatoyannopoulos, J.A. (2010). Sequencing newly replicated DNA reveals widespread plasticity in human replication timing. Proc. Natl. Acad. Sci. U. S. A.

Haradhvala, N.J., Polak, P., Stojanov, P., Covington, K.R., Shinbrot, E., Hess, J.M., Rheinbay, E., Kim, J., Maruvka, Y.E., Braunstein, L.Z., et al. (2016). Mutational Strand Asymmetries in Cancer Genomes Reveal Mechanisms of DNA Damage and Repair. Cell 164, 538–549.

Huang, X., Wojtowicz, D., and Przytycka, T.M. (2018). Detecting presence of mutational signatures in cancer with confidence. Bioinformatics 34, 330–337.

Hutchins, L.N., Murphy, S.M., Singh, P., and Graber, J.H. (2008). Position-dependent motif characterization using non-negative matrix factorization. Bioinformatics.

Hyka-Nouspikel, N., and Nouspikel, T. (2011). Nucleotide excision repair and B lymphoma: Somatic hypermutation is not the only culprit. Cell Cycle.

Jager, M., Blokzijl, F., Kuijk, E., Bertl, J., Vougioukalaki, M., Janssen, R., Besselink, N., Boymans, S., de Ligt, J., Pedersen, J.S., et al. (2019). Deficiency of nucleotide excision repair is associated with mutational signature observed in cancer. Genome Res. 1067–1077.

Jerbi, M., Ben Rekaya, M., Naouali, C., Jones, M., Messaoud, O., Tounsi, H., Nagara, M., Chargui, M., Kefi, R., Boussen, H., et al. (2016). Clinical, genealogical and molecular investigation of the xeroderma pigmentosum type C complementation group in Tunisia. Br. J. Dermatol. 174, 439–443.

Jolly, C., and Van Loo, P. (2018). Timing somatic events in the evolution of cancer. Genome Biol. 19, 1–9.

Kent, W.J., Sugnet, C.W., Furey, T.S., Roskin, K.M., Pringle, T.H., Zahler, A.M., and Haussler, a. D. (2002). The Human Genome Browser at UCSC. Genome Res.

Köster, J., and Rahmann, S. (2012). Snakemake-a scalable bioinformatics workflow engine. Bioinformatics.

Kraemer, K.H. (1987). Xeroderma pigmentosum. Cutaneous, ocular, and neurologic abnormalities in 830 published cases. Arch. Dermatol.

Kraemer, K.H., Lee, M.M., Andrews, A.D., and Lambert, W.C. (1994). The Role of Sunlight and DNA Repair in Melanoma and Nonmelanoma Skin Cancer: The Xeroderma Pigmentosum Paradigm. Arch. Dermatol.

Lehmann, A.R., McGibbon, D., and Stefanini, M. (2011). Xeroderma pigmentosum. Orphanet J. Rare Dis. 6, 1–6.

Li, H., and Durbin, R. (2009). Fast and accurate short read alignment with Burrows-Wheeler transform. Bioinformatics 25, 1754–1760.

Li, H., Handsaker, B., Wysoker, A., Fennell, T., Ruan, J., Homer, N., Marth, G., Abecasis, G., and Durbin, R. (2009). The Sequence Alignment/Map format and SAMtools. Bioinformatics 25, 2078–2079.

Ma, X., Liu, Y., Liu, Y., Alexandrov, L.B., Edmonson, M.N., Gawad, C., Zhou, X., Li, Y., Rusch, M.C., John, E., et al. (2018). Pan-cancer genome and transcriptome analyses of 1,699 paediatric leukaemias and solid tumours. Nature 555, 371–376.

Matsuda, T., Bebenek, K., Masutani, C., Rogozin, I.B., Hanaoka, F., and Kunkel, T.A. (2001). Error rate and specificity of human and murine DNA polymerase η. J. Mol. Biol.

Melis, J.P.M., Wijnhoven, S.W.P., Beems, R.B., Roodbergen, M., van den Berg, J., Moon, H., Friedberg, E., van der Horst, G.T.J., Hoeijmakers, J.H.J., Vijg, J., et al. (2008). Mouse Models for Xeroderma Pigmentosum Group A and Group C Show Divergent Cancer Phenotypes. Cancer Res. 68, 1347–1353.

Melis, J.P.M., Kuiper, R. V., Zwart, E., Robinson, J., Pennings, J.L.A., van Oostrom, C.T.M., Luijten, M., and Van Steeg, H. (2013). Slow accumulation of mutations in Xpc−/− mice upon induction of oxidative stress. DNA Repair (Amst).

Neph, S., Kuehn, M.S., Reynolds, A.P., Haugen, E., Thurman, R.E., Johnson, A.K., Rynes, E., Maurano, M.T., Vierstra, J., Thomas, S., et al. (2012). BEDOPS: High-performance genomic feature operations. Bioinformatics.

Oetjen, K.A., Levoska, M.A., Tamura, D., Ito, S., Douglas, D., Khan, S.G., Calvo, K.R., Kraemer, K.H., and DiGiovanna, J.J. (2019). Predisposition to hematologic malignancies in patients with xeroderma pigmentosum. Haematologica haematol.2019.223370.

Pedersen, B.S., and Quinlan, A.R. (2018). Mosdepth: Quick coverage calculation for genomes and exomes. Bioinformatics.

Petryk, N., Kahli, M., D’Aubenton-Carafa, Y., Jaszczyszyn, Y., Shen, Y., Silvain, M., Thermes, C., Chen, C.L., and Hyrien, O. (2016). Replication landscape of the human genome. Nat. Commun. 7, 1–13.

Quinlan, A.R. (2014). BEDTools: The Swiss-Army tool for genome feature analysis. Curr. Protoc. Bioinforma.

R Development Core Team, R. (2011). R: A Language and Environment for Statistical Computing.

Ramos, A.H., Lichtenstein, L., Gupta, M., Lawrence, M.S., Pugh, T.J., Saksena, G., Meyerson, M., and Getz, G. (2015). Oncotator: Cancer variant annotation tool. Hum. Mutat.

Robbe, P., Popitsch, N., Knight, S.J.L., Antoniou, P., Becq, J., He, M., Kanapin, A., Samsonova, A., Vavoulis, D. V., Ross, M.T., et al. (2018). Clinical whole-genome sequencing from routine formalin-fixed, paraffin-embedded specimens: pilot study for the 100,000 Genomes Project. Genet. Med.

Sarasin, A., Blanchet-Bardon, C., Renault, G., Lehmann, A., Arlett, C., and Dumez, Y. (1992). Prenatal diagnosis in a subset of trichothiodystrophy patients defective in DNA repair. Br. J. Dermatol.

Sarasin, A., Quentin, S., Droin, N., Sahbatou, M., Saada, V., Auger, N., Boursin, Y., Dessen, P., Raimbault, A., Asnafi, V., et al. (2019). Familial predisposition to TP53/complex karyotype MDS and leukemia in DNA repair-deficient xeroderma pigmentosum. Blood.

Schumann, F., Blanc, E., Messerschmidt, C., Blankenstein, T., Busse, A., and Beule, D. (2019). SigsPack, a package for cancer mutational signatures. BMC Bioinformatics 20, 1–9.

Seplyarskiy, V., Akkuratov, E.E., Akkuratova, N. V., Andrianova, M.A., Nikolaev, S.I., Bazykin, G.A., Adameyko, I., and Sunyaev, S.R. (2018). Error-prone bypass of DNA lesions during lagging strand replication is a common source of germline and cancer mutations. BioRxiv 200691.

Seplyarskiy, V.B., Soldatov, R.A., Popadin, K.Y., Antonarakis, S.E., Bazykin, G.A., and Nikolaev, S.I. (2016). APOBEC-induced mutations in human cancers are strongly enriched on the lagging DNA strand during replication. 1–9.

Sethi, M., Lehmann, A.R., Fawcett, H., Stefanini, M., Jaspers, N., Mullard, K., Turner, S., Robson, A., McGibbon, D., Sarkany, R., et al. (2013). Patients with xeroderma pigmentosum complementation groups C, e and v do not have abnormal sunburn reactions. Br. J. Dermatol. 169, 1279–1287.

Shen, R., and Seshan, V.E. (2016). FACETS: Allele-specific copy number and clonal heterogeneity analysis tool for high-throughput DNA sequencing. Nucleic Acids Res. 44.

Soufir, N., Ged, C., Bourillon, A., Austerlitz, F., Chemin, C., Stary, A., Armier, J., Pham, D., Khadir, K., Roume, J., et al. (2010). A Prevalent Mutation with Founder Effect in Xeroderma Pigmentosum Group C from North Africa. J. Invest. Dermatol. 130, 1537–1542.

Stone, J.E., Lujan, S.A., Kunkel, T.A., and Kunkel, T.A. (2012). DNA polymerase zeta generates clustered mutations during bypass of endogenous DNA lesions in Saccharomyces cerevisiae. Environ. Mol. Mutagen.

Thurman, R.E., Day, N., Noble, W.S., and Stamatoyannopoulos, J.A. (2007). Identification of higher-order functional domains in the human ENCODE regions. Genome Res.

Tomasetti (2016). Stem cell divisions, somatic mutations, cancer etiology, and cancer prevention. Int. Encycl. Public Heal. 80, 381–388.

Waszak, S.M., Northcott, P.A., Buchhalter, I., Robinson, G.W., Sutter, C., Groebner, S., Grund, K.B., Brugières, L., Jones, D.T.W., Pajtler, K.W., et al. (2018). Spectrum and prevalence of genetic predisposition in medulloblastoma: a retrospective genetic study and prospective validation in a clinical trial cohort. Lancet Oncol. 19, 785–798.

Wijnhoven, S.W.P., Kool, H.J.M., Mullenders, L.H.F., Van Zeeland, A.A., Friedberg, E.C., Van Der Horst, G.T.J., Van Steeg, H., and Vrieling, H. (2000). Age-dependent spontaneous mutagenesis in Xpc mice defective in nucleotide excision repair. Oncogene 19, 5034–5037.

Yoon, T., Chakrabortty, A., Franks, R., Valli, T., Kiyokawa, H., and Raychaudhuri, P. (2005). Tumor-prone phenotype of the DDB2-deficient mice. Oncogene 24, 469–478.

Yung, C.K., O’Connor, B.D., Yakneen, S., Zhang, J., Ellrott, K., Kleinheinz, K., Miyoshi, N., Raine, K.M., Royo, R., Saksena, G.B., et al. (2017). Large-Scale Uniform Analysis of Cancer Whole Genomes in Multiple Computing Environments. BioRxiv.

Zheng, C.L., Wang, N.J., Chung, J., Moslehi, H., Sanborn, J.Z., Hur, J.S., Collisson, E.A., Vemula, S.S., Naujokas, A., Chiotti, K.E., et al. (2014). Transcription Restores DNA Repair to Heterochromatin, Determining Regional Mutation Rates in Cancer Genomes. Cell Rep. 9, 1228–1234.

